# Synthetic carbon fixation via the autocatalytic serine threonine cycle

**DOI:** 10.1101/2022.09.28.509898

**Authors:** Sebastian Wenk, Vittorio Rainaldi, Hai He, Karin Schann, Madeleine Bouzon, Volker Döring, Steffen N. Lindner, Arren Bar-Even

**Affiliations:** Max Planck Institute of Molecular Plant Physiology, Am Mühlenberg 1, 14476 Potsdam-Golm, Germany; Max Planck Institute of Terrestrial Microbiology, Karl-von-Frisch-Str. 10, 35043 Marburg, Germany; Génomique Métabolique, Genoscope, Institut François Jacob, CEA, CNRS, Univ Evry, Université Paris-Saclay-4, 91057 Evry-Courcouronnes, France; Department of Biochemistry, Charité Universitätsmedizin, Virchowweg 6, 10117 Berlin, Germany

## Abstract

Atmospheric CO_2_ poses a major threat to life on Earth by causing global warming and climate change. On the other hand, it is the only carbon source that is scalable enough to establish a circular carbon economy. Accordingly, technologies to capture and convert CO_2_ to reduced one-carbon (C_1_) molecules (e.g. formate) using renewable energy are improving fast. Driven by the idea of creating sustainable bioproduction platforms, natural and synthetic C_1_-utilization pathways are engineered into industrially relevant microbes. The realization of synthetic C_1_-assimilation cycles in living organisms is a promising but challenging endeavour. Here, we engineer the autocatalytic serine threonine cycle, a synthetic C_1_-assimilation route in *Escherichia coli*. Our stepwise engineering approach in tailored selection strains combined with adaptive laboratory evolution experiments enabled the organism to grow on formate. The synthetic strain uses formate as the sole carbon and energy source and is capable of growing at ambient CO_2_ concentrations, demonstrating the feasibility of establishing synthetic C_1_-assimilation cycles over laboratory timescales.

## Introduction

The transition from a fossil-based, CO_2_-emitting economy towards a circular CO_2_-neutral economy is an imminent challenge to avert climate catastrophes^1^. CO_2_ is threatening our way of life by causing climate change and global warming, but, at the same time it is the only feedstock scalable enough to provide carbon for chemical or microbial production processes. Chemistry offers several efficient options to reduce CO_2_ into simple one-carbon (C_1_) molecules like formate or methanol, however, multi-carbon molecules cannot be produced with a high productivity_2,3_. Here, microbial cell factories can fill the gap to produce more complex value-added chemicals. Driven by this concept, industrially relevant model microbes have been engineered to grow on C_1_ molecules as sole carbon sources ^4–8^. Formate has been determined a key C_1_ molecule, as it can be produced very efficiently from CO_2_ and presents similar physicochemical properties to methanol ^9–12^. In nature, formate serves as a carbon and energy source to organisms growing via the serine cycle, the reductive glycine pathway (rGlyP) and the Calvin–Benson–Bassham (CBB) cycle (aerobic growth) or the reductive acetyl-CoA pathway (anaerobic growth)^2,13–16^. To facilitate the engineering of formatotrophy in industrially relevant microbes, several synthetic formate assimilation pathways were suggested and characterized computationally^2,17,18^. Among them, the serine threonine cycle (STC) was described as highly suitable for implementation in model microbes like *E. coli* and for biotechnological applications (Figure 1). It is oxygen tolerant, operates at ambient CO_2_ concentrations, requires the expression of only a few foreign enzymes and since it is an autocatalytic cycle, it has the potential to support exponential growth^19^. Autocatalytic cycles are very common in nature due to their optimal network topology but are considered difficult to engineer as they are susceptible to instabilities^20^. The design of the STC resembles the structure of the natural serine cycle (Supplementary Figure 1)^13^, but is modified to rely on the endogenous metabolism of *E. coli* for most of the pathway reactions^17^.

**Figure 1:**
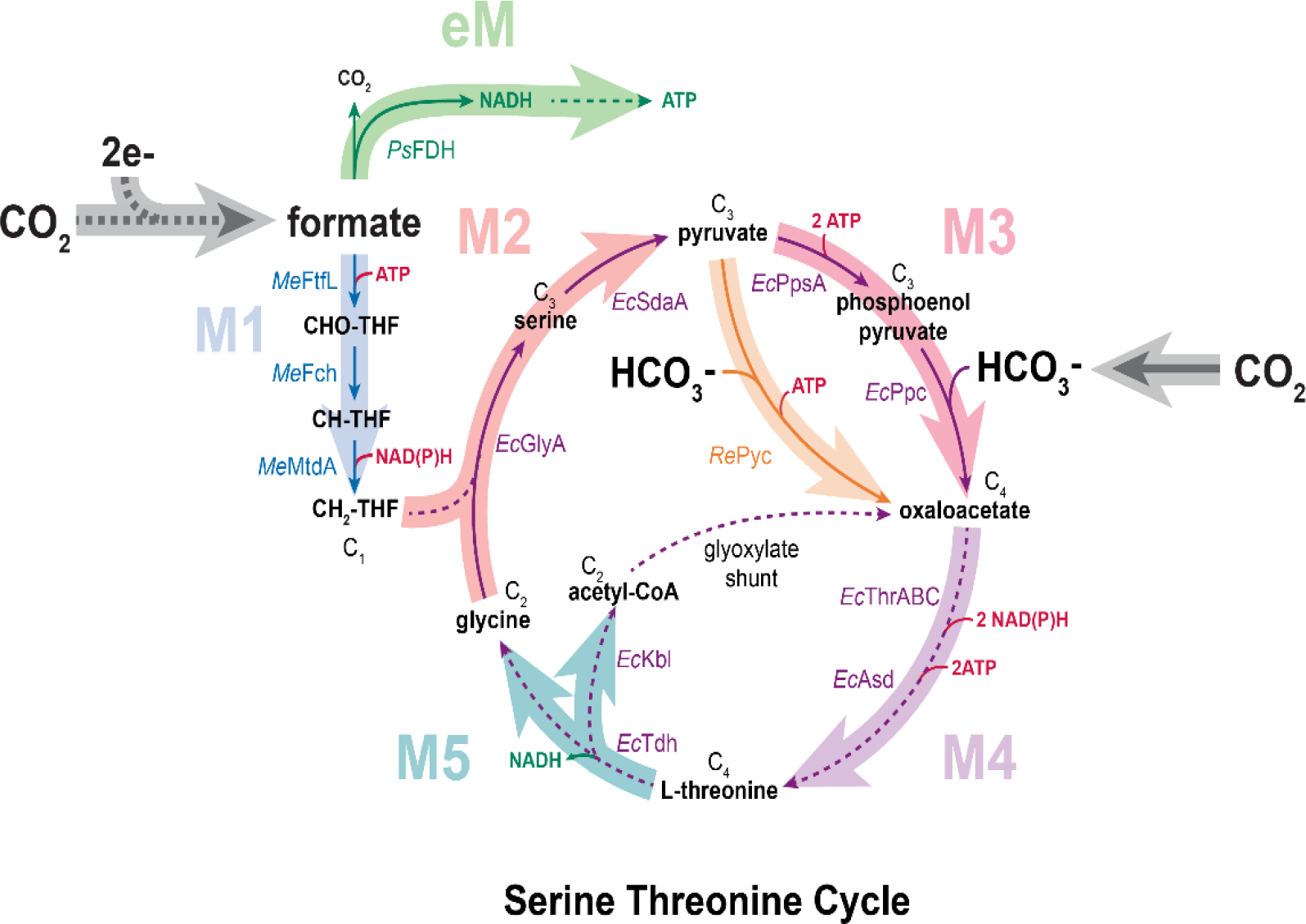
The autocatalytic serine threonine cycle is suited for synthetic carbon fixation in *E. coli*. To run the synthetic cycle in *E. coli*, most of the reactions can be catalyzed by endogenous enzymes (purple arrows). CO_2_-derived formate is converted into the one-carbon (C_1_) molecule methylene-THF (CH_2_-THF) via module 1 (M1) of the STC. The carbon from CH_2_-THF is attached to glycine producing the three-carbon (C_3_) molecule serine which is converted to pyruvate (M2). In M3 (the four-carbon (C_4_) molecule), pyruvate is converted to oxaloacetate fixing CO_2_ in the form of HCO_3-_ either via phosphoenolpyruvate (PEP) carboxylation (red arrow) or direct carboxylation of pyruvate (orange arrow). From oxaloacetate, threonine is produced (M4) and then cleaved into the two-carbon (C_2_) molecules glycine and acetyl-CoA (M5). As acetyl-CoA can be converted into oxaloacetate via the glyoxylate shunt, the cycle becomes autocatalytic. An energy module (eM) consisting of a NAD-dependent formate dehydrogenase supplies the cell with reduction equivalents and ATP derived from formate. CO_2_ reduction to formate is achieved electrochemically (grey dotted arrow). *Ps, Pseudomonas sp. (strain 101)*; *Me, Methylorubrum extorquens*; *Ec, Escherichia coli*; *Re, Rhizobium etli;* THF, tetrahydrofolate; Fdh, formate dehydrogenase; FtfL, formate-THF ligase; Fch, methenyl-THF cyclohydrolase; MtdA, methylene-THF dehydrogenase; GlyA, serine hydroxymethyltransferase; SdaA, serine deaminase; PpsA, PEP synthetase; Ppc, PEP carboxylase; Pyc, pyruvate carboxylase; ThrA, aspartate kinase I / homoserine dehydrogenase I; ThrB, homoserine kinase; ThrC, threonine synthase; Asd, aspartate-semialdehyde dehydrogenase; Tdh, threonine dehydrogenase and Kbl, 2-amino-3-ketobutyrate CoA ligase.

The implementation of the long and complex cycle in the heart of *E. coli’s* metabolism (the STC overlaps with glycolysis and the highly regulated PEP–pyruvate–oxaloacetate node ^21^) requires a substantial rerouting of central metabolic fluxes and is expected to reveal fundamental insights into the plasticity of the central metabolic network. In recent years, several studies enabled C_1_-dependent growth of *E. coli* via natural autocatalytic cycles, overlapping with the pentose phosphate pathway and glycolysis: autotrophic growth via the CBB cycle^4^ and methylotrophic growth via the ribulose monophosphate (RuMP) cycle^6,22^. Also, very recently a synthetic autocatalytic cycle for CO_2_ fixation was demonstrated in the strict anaerobe *Clostridium ljungdahlii*^23^. For formatotrophic growth, the non-autocatalytic, linear rGlyP has been established in *E. coli* ^5^, but previous attempts to establish a natural autocatalytic formate assimilation cycle^24^ or a synthetic formaldehyde assimilation cycle^25^ required the addition of co-substrates and so far, cyclic pathway activity was not reported. Thus, the engineering of the autocatalytic, new-to-nature STC in *E. coli* would overcome an open challenge to synthetic biology.

Here, we report the establishment of the complete STC and the conversion of the obligate heterotroph *E. coli* to a full formatotroph. We show that a combination of targeted engineering and adaptive laboratory evolution (ALE) enables the activity of the complete cycle. First, we engineered auxotrophic strains that depended on pathway activity for growth^26^. Then, we expressed individual pathway modules to complement the strain’
ss auxotrophies. Finally, we used ALE to overcome metabolic bottlenecks that prevented cyclic pathway activity achieving formatotrophic growth via the STC within ∼200 days of evolution. After confirming the formatotrophic phenotype of isolated strains by growth and isotopic labelling experiments, we analysed genomic alterations via genome sequencing and determined a small number of mutations that enabled activity of the STC. This study shows for the first time a fully functional autocatalytic and new-to-nature C_1_-assimilation cycle in *E. coli*, which enables the organism to grow on the sustainable C_1_-substrate formate at ambient CO_2_ concentration.

## Results

### Pathway modularization enables the stepwise engineering of the STC

To facilitate the engineering of the STC in *E. coli*, we divided the pathway into six metabolic modules (M) (Figure 1) that could be tested individually or in combination in dedicated selection strains: (M1) the C_1_ module that converts CO_2_-derived formate into CH_2_-THF; (M2) the C_3_ module that converts CH_2_-THF and glycine into pyruvate; (M3) the C_4_ module that converts pyruvate into oxaloacetate; (M4) the threonine synthesis module that drives the carbon flux from oxaloacetate towards threonine; (M5) the threonine cleavage module that cleaves threonine into glycine and acetyl-CoA and (eM) the energy module that oxidizes formate to generate NADH^27^. Module enzymes that are not naturally encoded in the *E. coli* genome were cloned into synthetic operons using defined promoters and ribosome binding sites (RBS) to modulate their expression levels and were expressed from the genome. Native *E. coli* enzymes were overexpressed when necessary by a genomic promoter exchange. Modules were tested for their *in vivo* activity in dedicated selection strains that depend on module activity for cell growth.

### Rewiring *E. coils* central metabolism for the implementation of the STC

As a substantial rewiring of *E. coli’s* central metabolism is required for the implementation of the STC - the main carbon flux needs to be directed from oxaloacetate to threonine (M4) which is cleaved to produce glycine and acetyl-CoA (M5) ^17^ - we first created a selection strain that depends on the combined activity of M4 and M5 for cell growth (hereafter referred to as *XGAS)*. This strain is practically auxotrophic to acetyl-CoA, serine and glycine as the canonical routes to these metabolites are blocked through gene deletions (Figure 2A). *XGAS* is only able to grow on glycerol if both M4 and M5 are sufficiently active to divert a major fraction of the cellular carbon flux via threonine synthesis and cleavage. When cultivated on glycerol, the strain was not able to grow, indicating that endogenous expression of M4 and M5 genes does not support sufficient flux towards acetyl-CoA and glycine. Only when the threonine cleavage module M5 was overexpressed from the genome (*XGAS:gM5*), growth on glycerol was observed (Figure 2B). Additional overexpression of M4 did not improve growth on glycerol, indicating that the creation of a strong sink for threonine through the expression of M5 also increased the flux towards its synthesis. We thus continued the engineering of the STC with the *XGAS:gM5*. In the next step, we included the formate assimilation module M1 into the pathway selection. To test for its activity, we deleted the glycine cleavage system (GCS) in the *XGAS:gM5* strain, creating *XGAS_C1::gM5*. The GCS deletion prevents glycine cleavage into CH_2_-THF and CO_2_. Thus, *XGAS_C1:gM5* is auxotrophic to metabolites derived from CH_2_-THF (e.g. methionine, purines, CoA and thymidine) and cannot produce serine from glycine. Expression of the enzymes of the C_1_-module (M1) in the *XGAS_C1:gM5* strain establishes formate as the precursor of the essential THF-bound C_1_ moieties. As previous studies determined M1 enzymes from *Methylorubrum extorquens* suitable for formate assimilation in *E. coli* ^5,28^, these genes were cloned into a synthetic operon and integrated into the genome of *XGAS_C1:gM5*. Upon integration of M1, the strain was able to grow on glycerol when formate was supplied as a co-substrate (Figure 2D).

**Figure 2:**
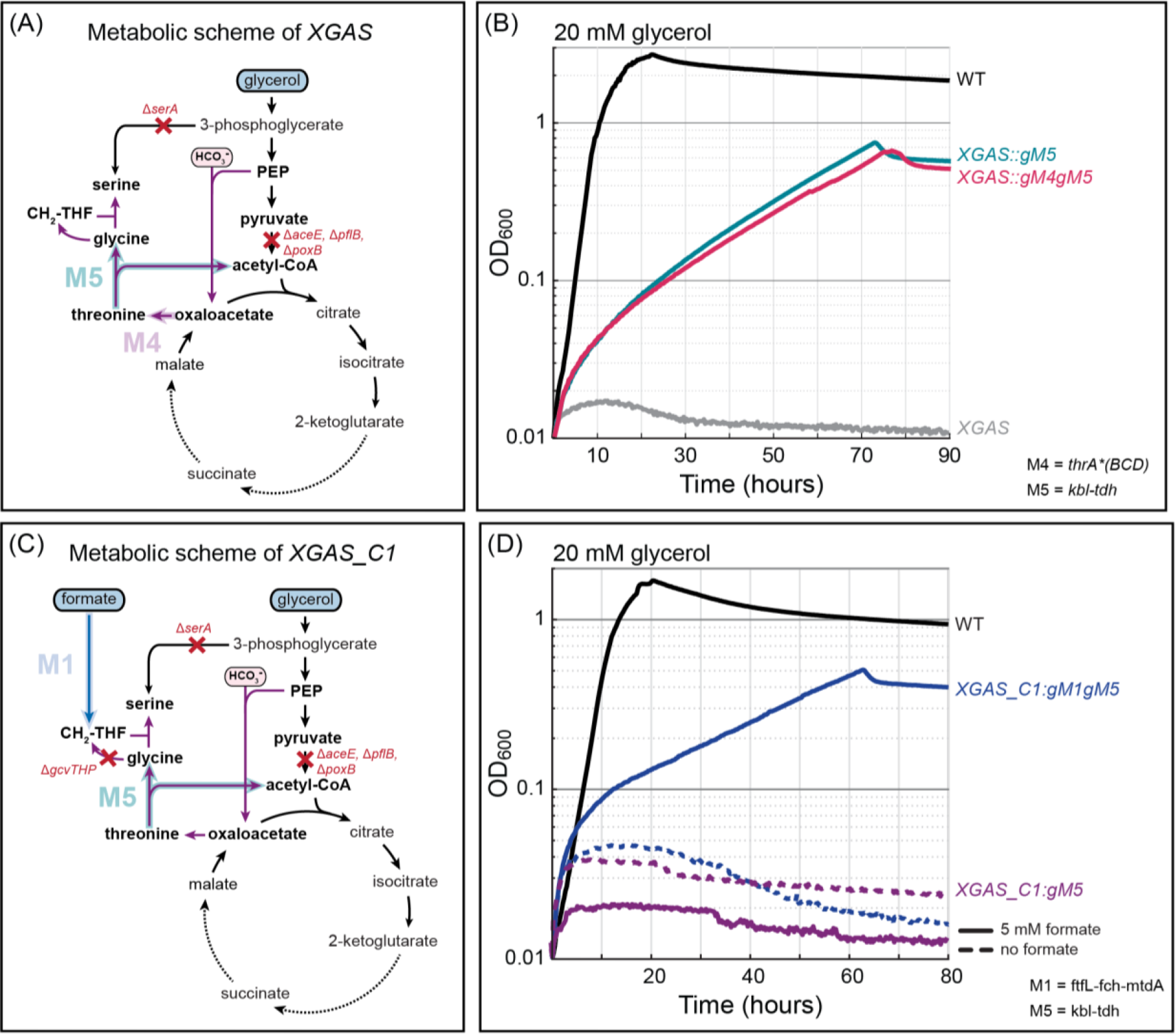
Rerouting of central metabolic fluxes enables growth via STC modules 1, 4 and 5. (A) Metabolic scheme of the *XGAS* selection strain that can only grow if modules 4 and 5 of the STC are sufficiently active to produce all cellular acetyl-CoA and glycine from threonine. (B) Overexpression of M5 from the genome (gM5) enabled the strain to grow on glycerol. Growth was not improved when also M4 was overexpressed from the genome. (C) Selection scheme of the strain *XGAS_C1* which cannot produce serine from glycine as the glycine cleavage system genes (*gcvTHP)* are deleted. The auxotrophy can only be released if formate is assimilated into CH_2_-THF by M1 enzymes. (D) When both M1 and M5 of the STC were expressed from the genome and formate was provided in the medium the strain was able to grow. Growth experiments were performed within 96-well plates in triplicates, which resulted in identical curves (±5%), and hence were averaged. All experiments (in triplicates) were repeated three times, which showed highly similar growth behavior. WT, wild type; *serA*, phosphoglycerate dehydrogenase; *aceE*, pyruvate dehydrogenase; *poxB*, pyruvate oxidase and *pflB*, pyruvate formate lyase.

### Optimization of pathway energetics and adaptive laboratory evolution enable cyclic pathway activity

The strain *XGAS_C1:gM1gM5* was able to grow on glycerol and formate via M1, M4 and M5 of the STC but could not grow on pyruvate and formate which also selects for the complete activity of M3 and is an important step towards complete STC activity (Supplementary Figure 2). The reason for that might be related to pathway energetics: the net energy investment for the conversion of pyruvate to acetyl-CoA and glycine via the STC (4 ATP and 1 NAD(P)H) is much higher compared to glycerol (2 ATP) while pyruvate is more oxidized than glycerol (energy of combustion of pyruvate ∼1160 kJ/mol and glycerol ∼1650 kJ/mol ^29^) (Figure 3A). Hence, it is likely that the cell cannot derive sufficient energy from pyruvate to run the pathway reactions.

**Figure 3:**
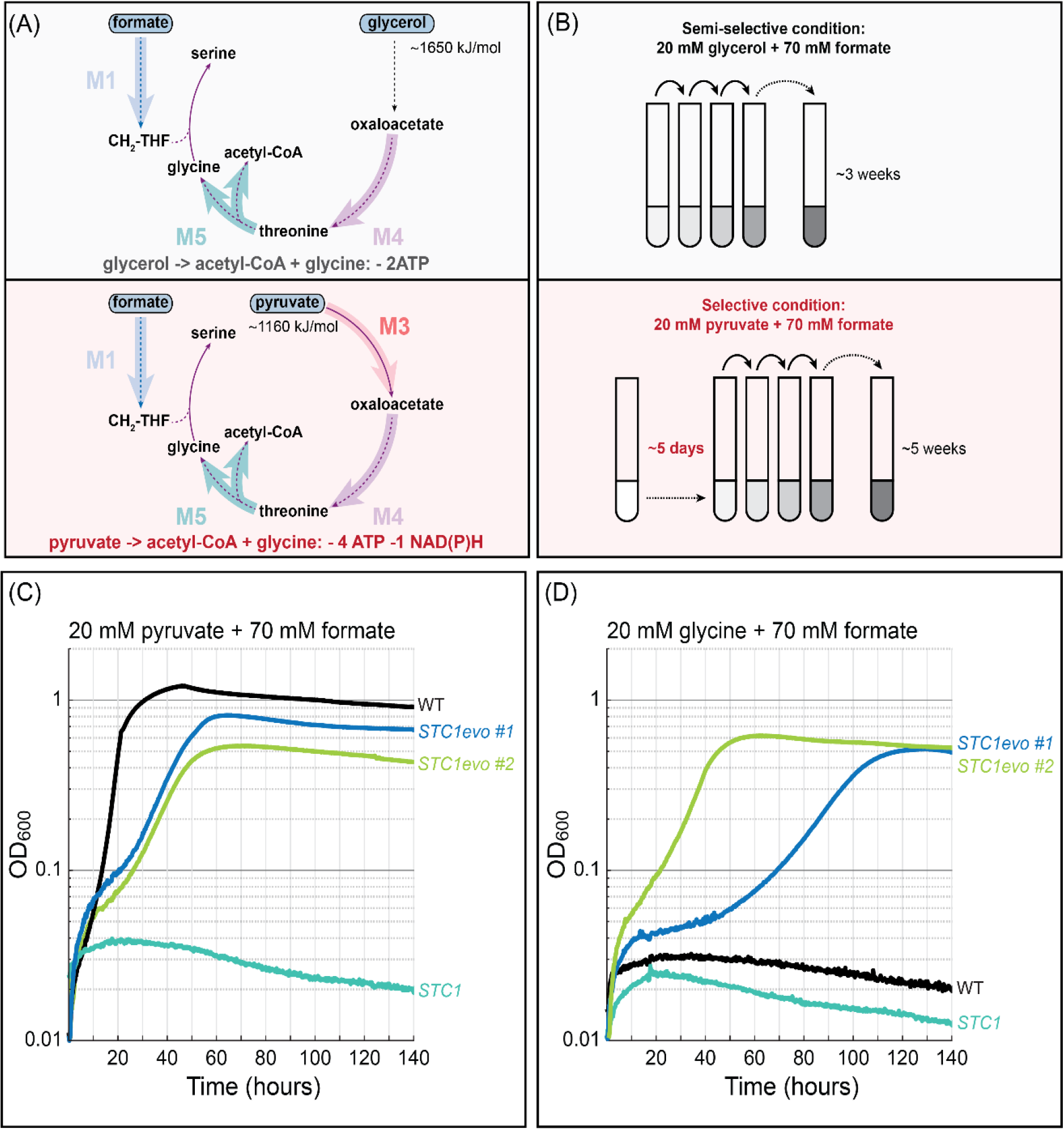
Strain optimization and adaptive laboratory evolution enable cyclic pathway activity (ALE1). (A) The conversion of pyruvate into acetyl-CoA and glycine via the STC requires the combined activity of M1, M3, M4 and M5. Furthermore, it requires the investment of additional two ATPs and one NAD(P)H compared to glycerol. This energetic difference might explain why *XGAS_C1:gM1gM5* was not able to grow on pyruvate and formate. (B) To achieve growth on pyruvate and formate, the a further engineered strain (*XGAS_C1:gM1gM2gM3gM5geM* (*STC1*)) was continuously cultivated in tubes with 20 mM glycerol and 70 mM formate for ~3 weeks thereby adapting the strain to high formate concentrations. Whenever the culture reached an OD_600_ > 0.5 it was diluted into fresh medium to an OD_600_ ~0.01. After ~3 weeks the medium was exchanged to fully selective medium containing 20 mM pyruvate and 70 mM formate. Approximately 5 days after the medium change the culture started to grow. Cultivation in selective medium was continued for ~5 weeks, then single colonies were isolated. (C) Isolated *STC1evo* strains were able to grow on pyruvate and formate and could also grow on (D) glycine and formate suggesting a fully active STC for the production of acetyl-CoA. Growth experiments were performed within 96-well plates in triplicates, which resulted in identical curves (±5%), and hence were averaged. All experiments (in triplicates) were repeated three times, which showed highly similar growth behavior. Abbreviations as in Figure 1.

To improve pathway energetics, we considered a pyruvate carboxylase (Pyc) as an alternative M3 and expression of the eM. The direct carboxylation of pyruvate to oxaloacetate shortens the STC by one reaction and reduces its ATP requirement by one ATP while the FDH of the eM supplies NADH from formate. After verifying Pyc activity *in vivo* (Supplementary Figure 3), we integrated *pyc* and the *fdh* into the genome of *XGAS_C1:gM1gM5*. As this strain was still not able to grow on pyruvate and formate, we concluded that other metabolic bottlenecks might exist that cannot be easily determined and addressed by rational engineering. Hence, we decided to use short-term ALE in tubes (ALE1) to enable sufficient pathway activity and achieve growth on pyruvate and formate (Figure 3B). We further engineered the strain to express M2 from the genome and conducted the ALE experiment as described in the methods section. After app. 9 weeks, we isolated single colonies capable of growing efficiently on pyruvate and formate via M1, M3, M4 and M5 of the STC (Figure 3C). Surprisingly, when testing the evolved strains with different carbon sources, we found that they were also able to grow on glycine and formate (Figure 3D). This indicated for the first time cyclic pathway activity of the STC and suggests that all modules are active *in vivo*: formate is assimilated into CH_2_-THF which is attached to glycine to form serine (M1 and M2). Then, serine is metabolized via the complete STC to produce acetyl-CoA required for cell growth (M3, M4 and M5).

### Developing formatotrophic growth via the STC

As complete activity of the STC was demonstrated for growth on glycine and formate, we aimed to achieve full formatotrophic growth via the pathway in the next step. This however requires at least a 1.63-fold increase in formate uptake (estimated by FBA; see Supplementary Figure 4 and methods) resulting in higher fluxes through the cycle and the energy module. As the strain *STC2* (an *STC1evo* strain with integrated *aceAB* operon) was not able to grow on formate only, we decided to evolve it in an automated setup (Chi.Bio) ^30^ under turbidostat conditions (ALE2), where glycine was provided as a limiting surrogate substrate which was stepwise decreased while the formate concentration was increased (Figure 4A). Starting from day ∼90, we completely omitted glycine in the growth medium. The sustained growth with formate as the sole carbon and energy source suggested the emergence and take-over of a glycine-independent strain that could grow formatotrophic. The cultivation with formate was continued for ∼60 days to improve the growth rate of the culture. Then, individual colonies were isolated and their formatotrophic growth was validated. The isolated evolved strains (*STC2evo* #1 and #2) grew robustly on formate without addition of another carbon source indicating that the STC was fully active (Figure 4B).

**Figure 4:**
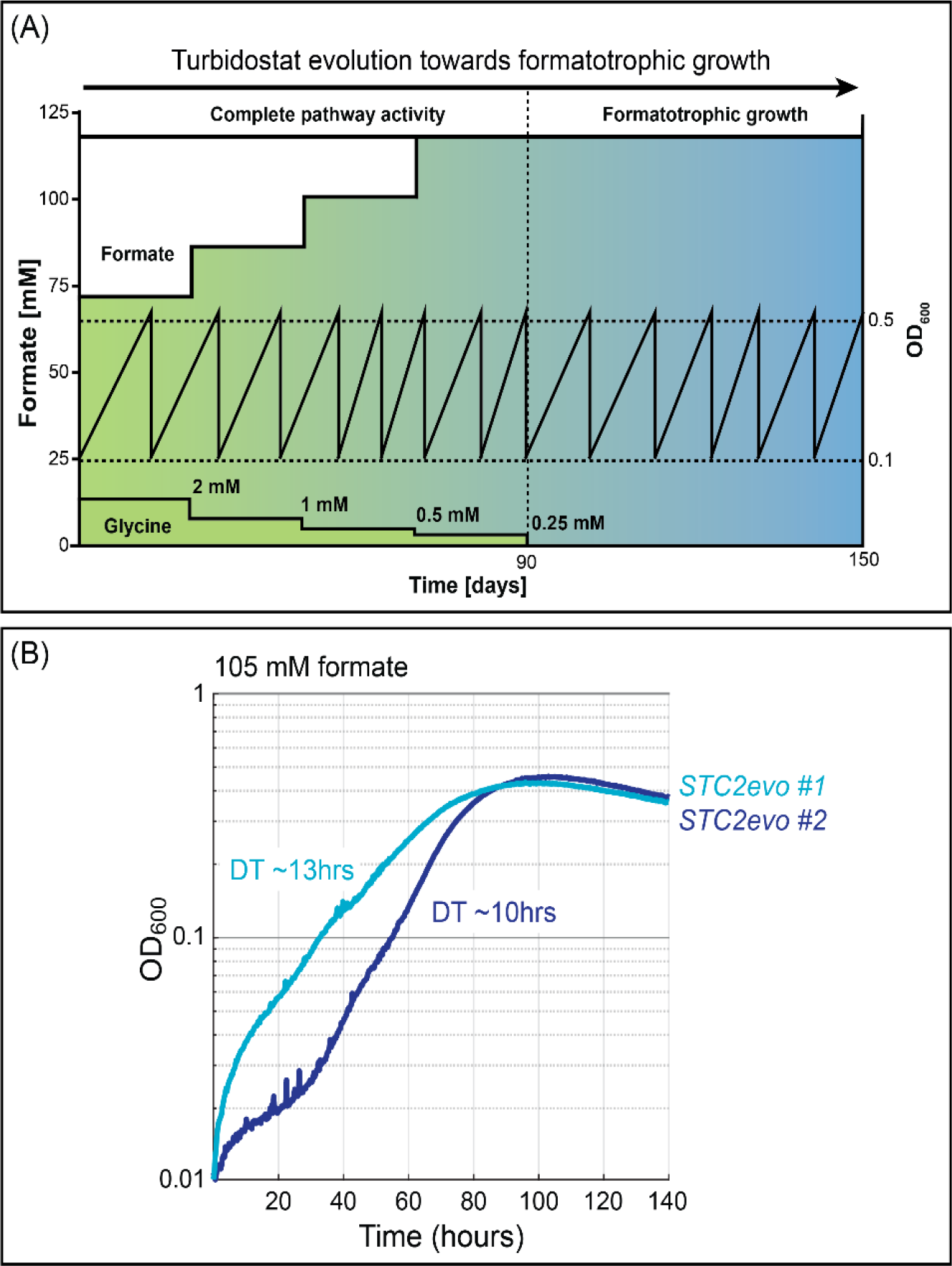
Automated adaptive laboratory evolution enables formatotrophic growth via the STC (ALE2). (A) Schematic of the automated adaptive laboratory evolution setup: In a Chi.Bio reactor the *STC1evo+* strain capable of growing on glycine and formate was continuously cultivated in a turbidostat mode keeping the culture in exponential phase (whenever the OD_600_ crossed the threshold of OD_600_ 0.5, the culture was diluted to OD_600_ 0.1 with fresh medium). Starting with a medium composition of 2 mM glycine and 70 mM formate, over the course of the experiment the glycine concentration was stepwise reduced to 0 mM while the formate concentration was increased to 120 mM. From day ~90 onwards, the culture started to grow fully formatotrophic. (B) Two isolated strains grew on formate only. Growth experiments were performed within 96-well plates in triplicates, which resulted in identical curves (±5%), and hence were averaged. All experiments (in triplicates) were repeated three times, which showed highly similar growth behavior. Doubling times (DT) are indicated in the figure.

### ^13^C isotopic labelling confirms formatotrophic growth via the STC

To confirm the incorporation of formate into *E. coli* metabolism, we performed a comprehensive isotopic labelling experiment. To this end, we cultivated the formatotrophic strain *STC2evo* on minimal medium supplemented with 105 mM ^13^C-formate and analysed the carbon labelling of proteinogenic amino acids. Feeding with ^13^C-formate should lead to incorporation of ^13^C into proteinogenic amino acids which can be detected by liquid chromatography–mass spectrometry (LC-MS). As expected, all pathway-related amino acids were labelled confirming their origin in formate and providing definitive evidence for the activity of the STC (Figure 5 and Supplementary Figure 5). The fact that some amino acids were not completely labelled can be explained by the incorporation of unlabelled HCO_3-_ via M3 of the STC (based on the labelling pattern of proline and arginine, we estimated ∼25% of intracellular CO_2_ to be unlabelled^4^) and the unlabelled fraction of the ^13^C-formate (99% pure).

**Figure 5:**
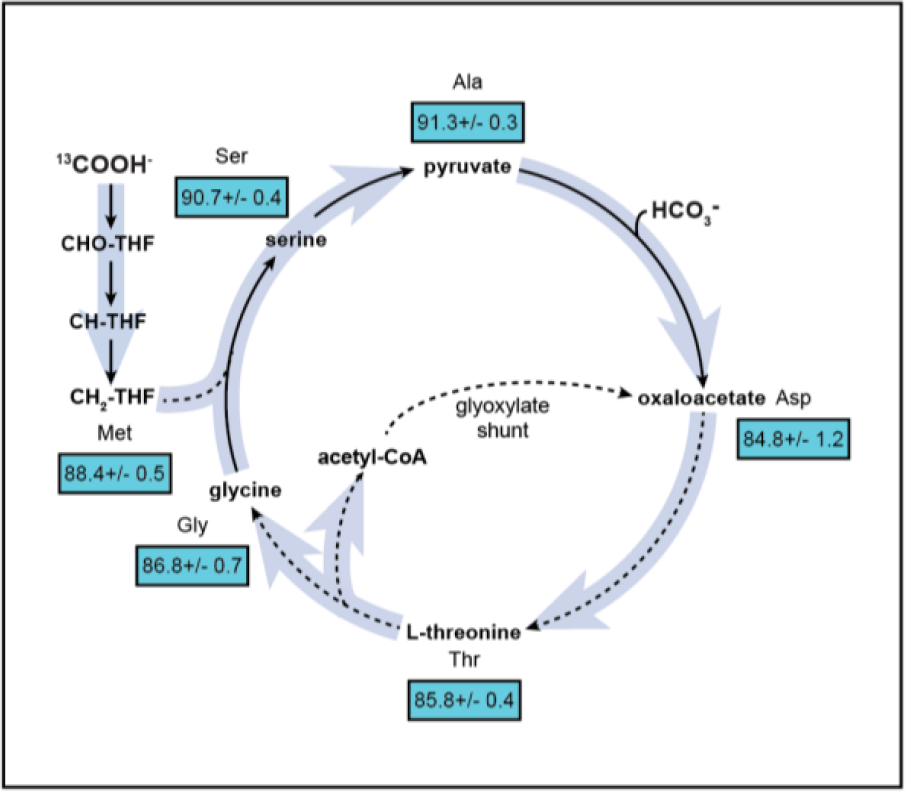
Incorporation of ^13^C-labelled formate confirms formatotrophic growth via the STC. The *STC2evo* strain capable of growing on formate was cultivated with 105 mM ^13^C-formate and its incorporation into proteinogenic amino acids was analysed via liquid chromatography–mass spectrometry. The analysis showed almost complete labelling of all proteinogenic amino acids. The numbers in the boxes represent the average total carbon labelling of the respective amino acid including standard deviation. The unlabelled fraction of amino acids is derived from unlabelled HCO_3-_assimilated via M3 of the STC. The experiment (in triplicates) was repeated three times and showed highly similar results.

### A small set of mutations enables formatotrophic growth via the STC

To elucidate the genetic adaptations that enabled cyclic pathway activity we sent genomic DNA of several candidate strains for genome sequencing. Apart from mutations that arose in individual strains and thus cannot be easily attributed to the pathway activity (Supplementary Table 2), all strains carried the same 39bp deletion between the leader peptide of the threonine biosynthesis operon (*thrL*) and the first gene of the operon (*thrA*) (Supplementary Figure 6). Analysing this genomic region in detail, we discovered that the deletion removes an RNA stem loop important for induction of attenuation in the presence of high intracellular threonine concentrations^31^. We assume that the 39bp deletion abolishes attenuation and enables constitutive expression of the of the *thrABC* operon even at elevated threonine levels occurring during growth via the STC. This hypothesis is supported by the fact that growth via the complete STC was abolished when we reconstituted the WT *thrLABC* operon in an evolved strain (Supplementary Figure 6). In addition to the 39bp deletion (which arose during ALE1), the formatotrophic strains incorporated a point mutation in the threonine biosynthesis gene *thrA* leading to a distinct amino acid exchange in the enzyme’s regulatory domain (S310P or S440P) which might influence the feedback inhibition of ThrA by threonine. Another mutation observed in all sequenced formatotrophic strains is a single base-pair substitution in the promoter region of the membrane-bound transhydrogenase (*pntAB*) that catalyses the proton transfer from NADH to NADP_+_. Surprisingly, the same mutation was found in a previous study where *E. coli* was engineered to grow on formate via the rGlyP_5_. In this study, quantitative PCR determined that the mutation increases transcript levels of *pntAB* by 13-fold. The higher expression of the transhydrogenase could have a beneficial effect on NAPDH availability, a key cofactor for the activity of the STC. While several other mutations were observed in the different evolved strains, the fact that they are not shared among all strains suggests that they are not strictly needed for optimized pathway activity. Hence it seems that a small number of mutations that influences expression and activity of M4 as well as pathway energetics is responsible for the activation of the STC in the engineered strains.

## Discussion

In this study, a new-to-nature autocatalytic formate assimilation cycle in a model microbe has been established. Using a strategy that couples pathway activity to the growth of the host organism combined with ALE, we showed that it is possible to convert the obligate heterotrophic organism *E. coli* into a full formatotroph over laboratory timescales. Our findings are in line with recent studies that successfully established natural C_1_ assimilation pathways in *E. coli* (CBB, rGlyP and RuMP) by similar strategies ^4–6,22^.

Compared to the natural serine cycle, the STC circumvents the emergence of the toxic pathway intermediate hydroxypyruvate and is harmonized to *E. coli’s* native metabolic set-up, avoiding the use of malyl-CoA synthetase and lyase which are absent in *E. coli’s* genome and could possibly interfere with the natural flux of metabolites through the TCA cycle when heterologously expressed. The STC is thus optimally suited for implementation in *E. coli* and other industrial relevant microbes with a similar central metabolism. Compared to other C_1_-assimilation routes, the autocatalytic STC shows the clear advantage of enabling formatotrophic growth at ambient CO_2_ which allows a wider application spectrum and easier cultivation conditions. Furthermore, a recent theoretical analysis showed that serine cycle variants can support formatotrophic growth with high energetic efficiencies^11^. While in this study energy and carbon metabolism were coupled, the energy module of the STC could be replaced by using an efficient methanol dehydrogenase with methanol as an auxiliary energy and possible carbon source.

The results of this study clearly highlight the importance of ALE for achieving pathway activity. While rational engineering provides the genetic setup for novel pathways and might suffice to enable pathway activity in some cases, it can hardly address the genetic fine tuning which is often required to balance fluxes between native and non-native metabolic reactions. This seems especially true for metabolic engineering approaches that aim to achieve novel growth modes of the host ^4–6,8^. In the case of the STC, ALE was used twice: first to enable cyclic pathway activity (ALE1, emergence of a Δ39bp deletion upstream of *thrABC*) and second to increase fluxes through the STC for achieving formatotrophic growth (ALE2, emergence of mutations in *thrA* and *pntAB*). Interestingly, only two evolutionary modifications can be directly linked to the pathway: (1) changes in the threonine synthesis operon, the only pathway module that was not rationally engineered in the strain and (2) changes in the expression of the membrane bound transhydrogenase. In contrast to what was observed for the engineering of natural autocatalytic C_1_-assimilation cycles (CBB^4^ and RuMP^6^), no tuning of metabolic branch points seem to be required to allow full growth via the STC. Looking at the study from the “Design, Build, Test, Learn” perspective of synthetic biology^32^ we can “learn” from these results that all pathway modules should have been overexpressed by engineering. Also, the transhydrogenase seems to be an important candidate gene for rational modifications, as the same mutation occurred independently in this and previous study^5^. Hence, future studies aiming to establish C_1_-assimilation routes requiring NADPH while using a NAD-dependent enzyme for energy generation should consider the overexpression of the transhydrogenase *ab initio*.

The results of this study clearly demonstrate the flexibility of *E. coli’s* central metabolic network and show for the first time that it is possible to introduce synthetic formate assimilation cycles in the heart of metabolism which enables robust formatotrophic growth. To create a platform strain for industrial applications, the formatotrophic strain created in this study could be combined with existing production modules to generate value-added chemicals from renewably produced formate ^33^. This provides a new option to develop production processes based on microbial cell factories growing on CO_2_-derived C_1_ substrates. Our study thus extends the solution space for designing processes based on CO_2_-derived feedstocks and points the way towards carbon negative chemical production in the framework of a circular economy.

## Supporting information

Gene Sequences

## Acknowledgements

The authors thank Lothar Willmitzer, Nico Claassens and Enrico Orsi for critical reading of the manuscript and helpful suggestions and Änne Michaelis for support with LC-MS measurements. SW thanks Caroline Gutjahr for guidance during the preparation of the manuscript. This work was funded by the Max Planck Society, by the German Ministry of Education and Research grant FormatPlant (part of BioEconomy 2030, Plant Breeding Research for the Bioeconomy), and MetAFor (031B0850B).

## Author contributions

A.B.-E. designed and supervised the research;

S.W. designed and conducted the experiments, analyzed the data and wrote the manuscript.

S.W. and K.S. genetically engineered *E. coli* for growth on formate

V.R. performed evolution experiments in Chi.Bio reactors, characterized isolated candidates and analyzed the experimental data

H.H. performed flux balance analysis

M.B. and V.D. designed and supervised evolution experiments

S.N.L. performed initial experiments and assisted in writing the manuscript

## Methods

### Chemicals and reagents

Primers and oligonucleotides were synthesized by Integrated DNA Technologies (IDT). PCR reactions were carried out either using DreamTaq polymerase or Phusion High-Fidelity DNA Polymerase (Thermo Fisher Scientific). Restriction digests and ligations were performed using FastDigest enzymes and T4 DNA ligase (Thermo Fisher Scientific). Glycerol, sodium pyruvate, sodium acetate, glycine, sodium formate and sodium formate-^13^C were ordered from Sigma-Aldrich.

### Strains

The *E. coli* strain SIJ488 was used for engineering purposes. *E. coli* strain DH5α and *E. coli* strain ST18 were used for cloning and conjugation purposes. All strains used in this study are listed in Supplementary Table 1.

### Strain engineering

Engineered strains were generated with recombineering techniques using SIJ488 as a base strain. For deletions, a linear dsDNA fragment containing an antibiotic resistance gene (kanamycin or chloramphenicol) was amplified with primers containing 50 bp homology arms flanking the target sequence to be deleted. The linear cassette was introduced into the strain via electroporation after induction of the recombination machinery with 15 mM arabinose for one hour. Colonies carrying the deletion were selected on LB plates supplemented with antibiotic. A similar procedure was used for genomic promoter exchange. To enable genomic overexpression of a synthetic operon, a conjugation-based genetic recombination method was used as described in^34^. In brief, the synthetic operon was cloned into a vector containing two 600bp homology regions compatible with the integration locus, chloramphenicol resistance gene (cam_R_), a levansucrase gene (sacB), and the conjugation gene traJI for the transfer of the plasmid. The resulting plasmid was transformed into chemically competent *E. coli* ST18 strains. Positive clones growing on chloramphenicol medium supplemented with 5-aminolevulinic acid (50 ug ml^-1^) were identified by colony PCR, and the confirmed recombinant ST18 strain was used as donor strain for the conjugation. Chloramphenicol resisting recipient *E. coli* strains were screened as positive strains for the first round of recombination. Subsequently, sucrose counter selection and kanamycin resistance tests were carried out to isolate recombinant *E. coli* strains with the correct synthetic operon integration into chromosome. All constructs were verified via PCR and sequencing.

### Synthetic-Operon construction

Before cloning the genes into synthetic operons, each gene was codon optimized for *E. coli* K-12 and an N-terminal 6xHis-tag was added. All genes were inserted into a cloning vector that attached a synthetic ribosome binding site upstream of the gene^35^. The same entry vector was then used for stepwise assembly of multi gene operons as described in^34^. Plasmids were constructed via restriction and ligation using standard kits and protocols supplied by Thermo Fischer Scientific. The assembled construct was then transferred to an expression vector containing a promoter and terminator sequence^34^.

### Growth media

Cloning and engineering steps were performed using Lysogeny Broth medium (10 g/l tryptone, 10 g/l NaCl, 5 g/l yeast extract + 15 g/l agar for plates) with the appropriate amount of relevant antibiotics (ampicillin/carbenicillin (100 ug/ml), kanamycin (50 ug/ml), chloramphenicol (30 ug/ml) and/or streptomycin (50 ug/ml)). Engineered and evolved strains were grown either in M9 minimal medium (50 mM Na_2_HPO_4_,20 mM KH_2_PO_4_, 1 mM NaCl, 20 mM NH_4_Cl, 2 mM MgSO_4_, and 100 μM CaCl_2_) or HEPES minimal medium (HMM, 200 mM HEPES, 1.32 mM K_2_HPO_4_, 0.5% NaCl, 1% NH_4_Cl, 2 mM MgSO_4_, and 100 μM CaCl_2_) supplemented with trace elements (50 mg/L EDTA, 31 mM FeCl_3_, 6.2 mM ZnCl_2_, 0.76 mM CuCl_2_-2H_2_O, 0.42 mM CoCl_2_-6H_2_O, 1.62 mMH_3_BO_3_, 81nM MnCl_2_-4H_2_O) and the relevant carbon sources.

### Growth experiments

For the cell growth tests, overnight cultures in LB medium were used to inoculate a pre-culture at an optical density (600 nm, OD_600_) of 0.02 in 4 ml fresh minimal medium under relaxing conditions (**as indicated in Supplementary Table 1**) in 14 ml glass test tubes. Cells were then cultivated in a shaking incubator at 37 °C and 220 rpm overnight. Cell cultures were harvested by centrifugation (11000 x g, 1 min), washed twice with fresh medium and used to inoculate the main culture, conducted aerobically either in 14 ml glass tube or Nunc 96-well microplates (Thermo Fisher Scientific) with appropriate carbon sources as indicated in the main text. In the 96 well plate cultivation, each well containing 150 μl culture was overlaid with 50 μl mineral oil (Sigma-Aldrich) to avoid evaporation while still allowing gas O_2_ and CO_2_ diffusion. Growth experiments were carried out in a BioTek Epoch2 plate reader (BioTek Instrument, USA) at 37 °C. OD_600_ was measured after a kinetic cycle of 12 shaking steps, which alternated between linear and orbital (1 mm amplitude), and were each 60 s long. OD_600_ values measured in the plate reader were calibrated to represent OD_600_ values in standard cuvettes according to OD_cuvette_ =OD_plate_/0.23. Glass tube culture was carried out in 4 ml of working volume, at 37 °C and shaking of 220 rpm. All growth experiments were performed in triplicate, and the growth curves shown represent the average of these triplicates.

### Tube evolution (ALE1)

The experiment was conducted with the STC1 strain and consisted of two consecutive steps: (1) cultivation under semi-selective conditions: glycerol + formate (selecting for the activity of M1, M4 and M5) and (2) cultivation under fully selective conditions: pyruvate + formate (selecting for the activity of M1, M3, M4 and M5) (Figure 3B). Several 14 ml glass tubes with biological replicates were cultivated in semi-selective medium (M9 minimal medium with 20 mM glycerol and 70 mM Na-formate) for 3 weeks. Whenever the strain reached an OD_600_ > 0.5 the culture was diluted into fresh medium to an OD_600_ of 0.01. After 3 weeks, the medium was changed to fully selective medium (20 mM pyruvate and 70 mM Na-formate). Five days after the medium swap, growth on pyruvate and formate was observed for the first time indicating the emergence of an evolved strain. Cultivation under selective conditions was continued for 5 weeks to further improve growth characteristics. Then, single colonies were isolated (*STC1evo*) and their growth on pyruvate and formate as well as glycine and formate was characterized as indicated in the main text.

### Turbidostat evolution experiment (ALE2)

The turbidostat evolution was performed using the automated Chi.Bio platform, set to dilute the cultures whenever the OD_600_ reached a set value equivalent to a cuvette OD_600_ of 0.5 in order to always keep the cells in exponential phase (avoiding mutations related to stationary phase adaptation) and to ensure a frequent medium turnover to avoid a detrimental increase in pH. The evolution was started with HMM200 (pH 7.2) containing 70 mM formate and 2 mM glycine supplemented with 50 ug/ml kanamycin sulfate to prevent contamination. Every ∼12 doublings (as estimated by the medium consumption rate), new medium was prepared halving the concentration of glycine. After several iterations (corresponding to 2 mM, 1 mM, 0.5 mM and 0.25 mM glycine), strains were isolated and tested for growth on formate without supplementation of glycine.

### ^13^C labeling experiment

For the labeling experiment the strain was inoculated from a plate into 3 ml LB with the appropriate antibiotic and grown overnight at 37 C at 250 rpm. The overnight culture was washed twice with minimal medium and a dilution was used to inoculate 4 ml minimal test medium containing ^13^C-labelled formate with an initial OD_600_ of 0.01. Upon reaching an OD_600_ of >0.5, was harvested for amino acid extraction.

### Sample preparation for LC-MS analysis

After harvesting the biomass, culture samples were prepared and analyzed as described previously^34^. In brief, for protein-bound amino acids, the equivalent of 1 ml culture at OD_600 ∼_1 were pelleted by centrifugation for 1 minute at 13,000 g. The resulting pellet was suspended in 1 mL of 6M HCl and incubated for 24 hours at 98C. The acid was then left to evaporate under a nitrogen stream, resulting in a dry hydrolysate. Dry hydrolysates were resuspended in 1 mL MilliQ water and centrifuged for 5 minutes at 14,000 g. The supernatant was then used for the LC-MS analysis. Hydrolyzed amino acids were separated using ultra performance liquid chromatography (Acquity, Waters, Milford, MA, USA) using a C18-reversed-phase column (Waters). Mass spectra were acquired using an Exactive mass spectrometer (Thermo Fisher). Data analysis was performed using Xcalibur (Thermo Fisher). Prior to analysis, amino-acid standards (Sigma-Aldrich) were analyzed under the same conditions in order to determine typical retention times.

### Whole-genome sequencing

Genomic DNA was extracted using a commercial kit (Mackerey Nagel) starting from 2-4 ml of overnight culture in LB. A minimum of 300 ng was sent for sequencing to NovoGene. Results were analyzed using the open source breseq software^36^. All NGS data was deposited at the Genome Sequence Archive.

### Flux balance analysis

Flux balance analysis (FBA) was used for comparing formate dependency of the STC under formate only or formate-glycine conditions. The modeling was conducted with COBRApy^37^ using the most updated *E. coli* genome-scale metabolic model *i*ML1515^38^ with curations and changes: (i) transhydrogenase (THD2pp) translocates one proton instead of two^39^; (ii) homoserine dehydrogenase (HSDy) was set to irreversibly produce homoserine^25^; (iii) anaerobic relevant reactions, PFL, OBTFL, FDR2, and FDR3, were removed from the model; (iv) POR5, GLYCK, FDH4pp FDH5pp, GART, DRPA, PAI2T, G6PDH2r, and ETHAAL were also knocked out to block unrealistic routes; (v) formate dehydrogenase (FDH) and pyruvate carboxylase (PYC) were implemented in the model; (vi) gene knock outs of the XGAS_C1 strain, i.e. *gcvTHF, aceE, poxB, serA* and additional *aceA* in the case of formate and glycine feeding condition, were also introduced in the model. We further fixed the upper and lower bounds of the biomass reaction and change the objective function to formate uptake (EX_for_e) to find the flux distribution resulting in biomass flux of the fixed value at the lowest possible flux through formate uptake, using formate only or formate and glycine as carbon sources. We expressed the formate dependency as the slope of biomass reaction, i.e. growth rate, and formate uptake rate from the modeling. The full code, including the changes described above, was deposited at https://github.com/he-hai/PubSuppl, within “2022_STC” the directory.

## Data availability

Additional information on the experimental setup as well as detailed results are available from the corresponding author upon request. Any strains and plasmids generated during this study are available upon completing a Materials Transfer Agreement.

## Code availability

MATLAB and breseq codes used for the analysis of the experiments are available from the corresponding author upon request.

