## Supplementary Figures for "Synthetic carbon fixation via the autocatalytic serine threonine cycle"

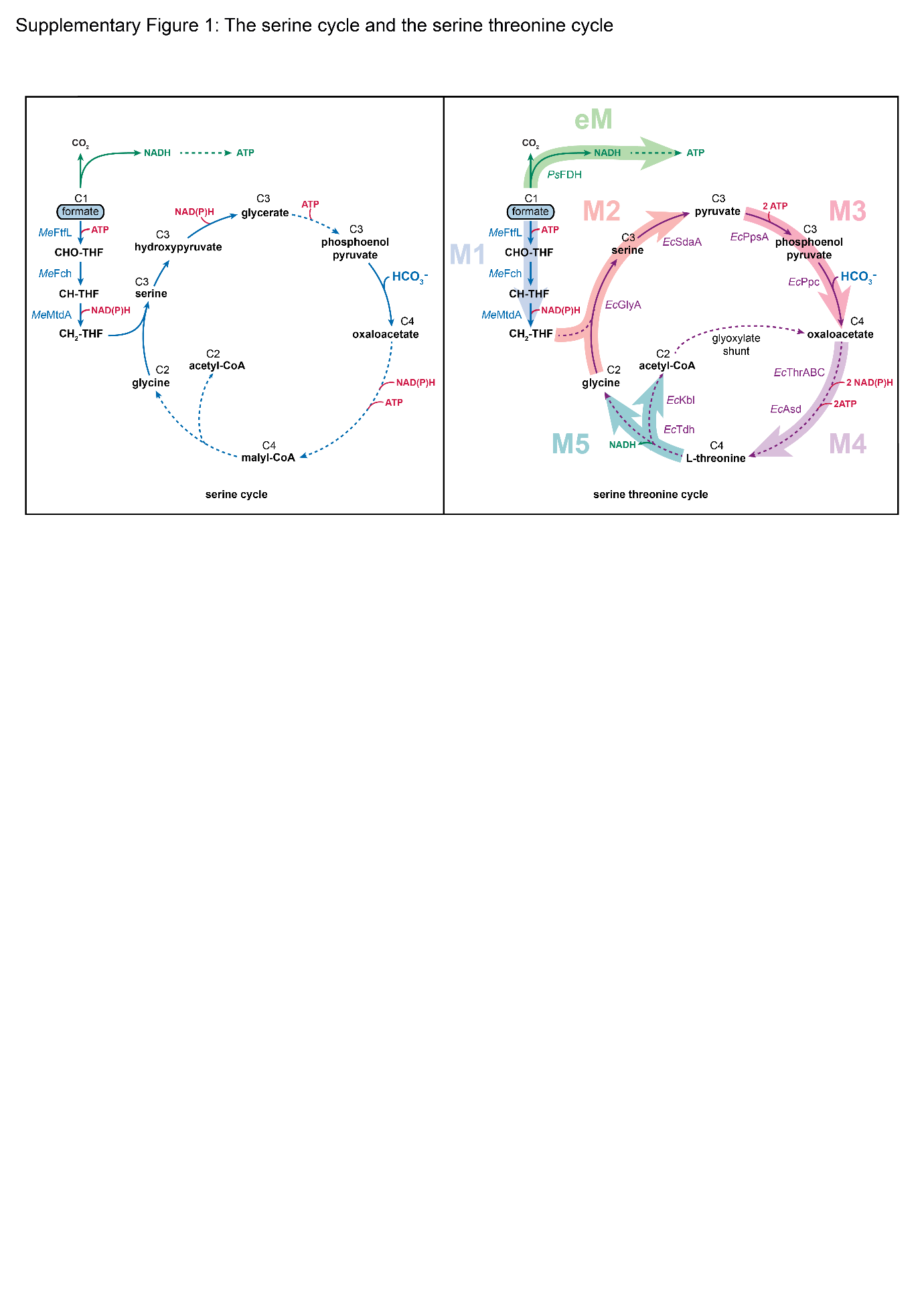

**Supplementary Figure 1: The serine threonine cycle follows the design principles of the natural serine cycle.**

(A) The design of the STC resembles the structure of the natural serine cycle: glycine + formate → serine → PEP + CO_2_ → C_4_ → glycine + acetyl-CoA, but is modified to rely on the endogenous metabolism of *E. coli* for most of the pathway reactions: (B) formate is assimilated by the enzyme FTL, which attaches it to the universal C_1_ carrier tetrahydrofolate (THF), forming 10-formyl-THF (CHO-THF). Then, CHO-THF is reduced to 5,10-methylene-THF (CH_2_-THF), which donates its formaldehyde moiety to glycine, giving rise to serine. In two enzymatic steps, serine is converted to PEP, bypassing the toxic intermediate hydroxypyruvate present in the natural serine cycle. PEP is then carboxylated to oxaloacetate. From oxaloacetate, the flux is directed to threonine, which is cleaved to recover glycine and generate acetyl-CoA as the pathway product. Ps, *Pseudomonas sp.* (strain 101); Me, *Methylorubrum extorquens*; Ec, *Escherichia coli*; THF, Tetrahydrofolate; Fdh, formate dehydrogenase; FtfL, formate-THF ligase; Fch, methenyl-THF cyclohydrolase; MtdA, methylene-THF dehydrogenase; GlyA, serine hydroxymethyltransferase; SdaA, serine deaminase; PEP, phosphoenolpyruvate; PpsA, PEP synthetase; Ppc, PEP carboxylase; Pyc, pyruvate carboxylase; ThrA, aspartate kinase I / homoserine dehydrogenase I; ThrB, homoserine kinase; ThrC, threonine synthase; Asd, aspartate-semialdehyde dehydrogenase; Tdh, threonine dehydrogenase and Kbl, 2-amino-3-ketobutyrate CoA ligase.

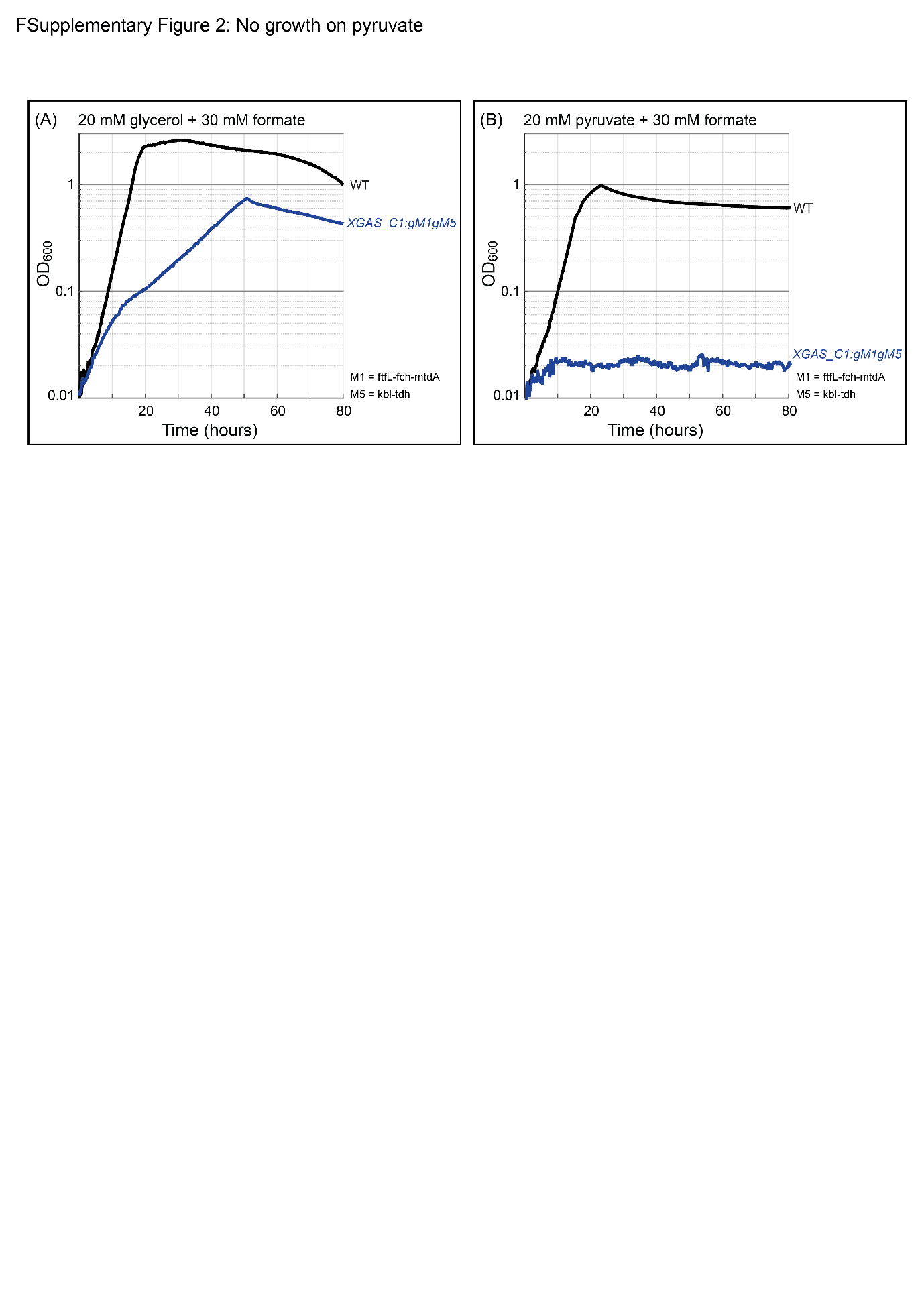

**Supplementary Figure 2: *XGAS_C1:gM1gM5* cannot grow on pyruvate and formate.**

(A) Genomic expression of M1 and M5 genes enabled the strain *XGAS_C1* to grow on glycerol when formate was supplemented. (B) No growth was observed when the canonical carbon source glycerol was replaced with pyruvate. Experiments were conducted within 96-well plates and were performed in triplicates, which displayed identical growth curves (±5%), and hence were averaged. All experiments (in triplicates) were repeated three times, which showed highly similar growth behavior. WT, wild type.

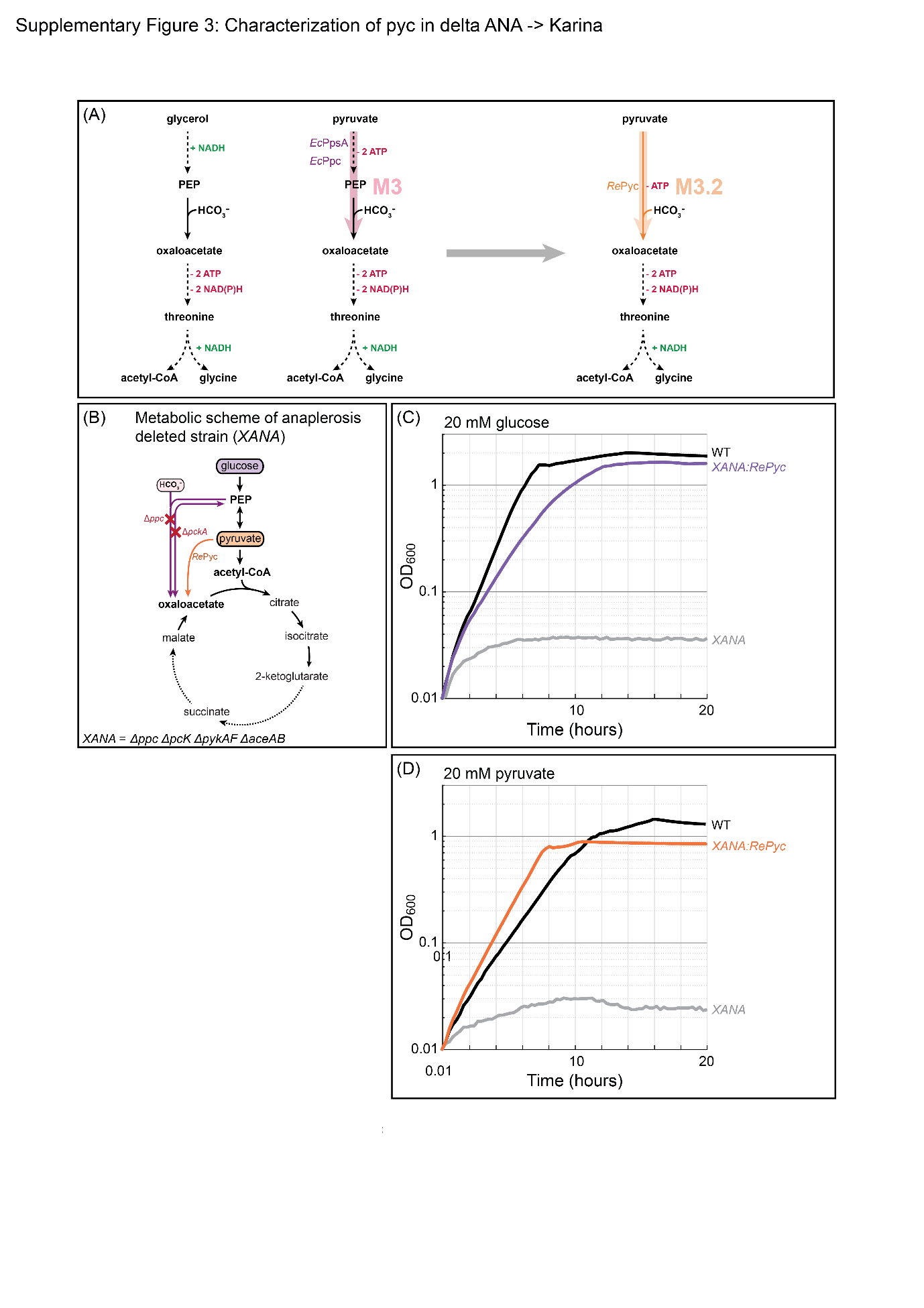

**Supplementary Figure 3: Implementation of an alternative module 3 improves pathway energetics of the STC.**

(A) To improve pathway energetics, we replaced the original M3 (*Ec*PpsA, EcPpc) with an alternative (M3.2) that requires only one enzymatic reaction and consumes only one ATP. To this end, we chose the pyruvate carboxylase from *Rhizobium etli* (*Re*Pyc). (B) Using a selection strain deleted in the anaplerotic reactions (Δ*ppc*Δ*pckA*) we tested the *in vivo* activity of *Re*Pyc. Only when the enzyme was expressed from the genome the strain was able to grow on (C) glycose and (D) pyruvate indicating the activity of *Re*Pyc and its suitability for the engineering of the STC. All experiments (in triplicates) were repeated three times, which showed highly similar growth behavior. WT, wild type.

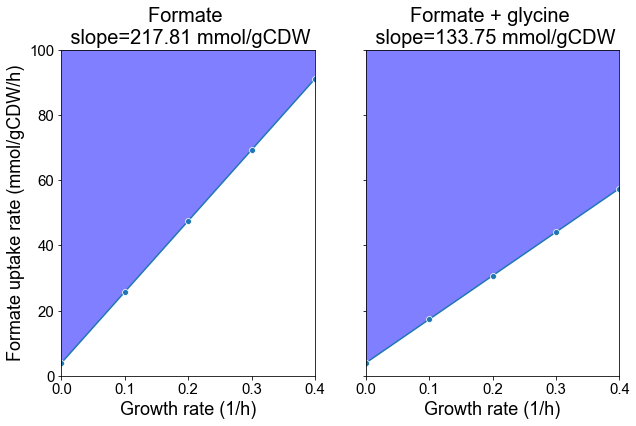

**Supplementary Figure 4: FBA estimation of formate dependency for cyclic STC activity.**

FBA analysis was conducted using a modified *E. coli* genome-scale model with the STC reactions and relevant genome manipulations. Minimal required formate uptake rates were calculated in different growth rates. The formate dependency was expressed as the slope between growth rates and formate uptake rates. See details in the methods section.

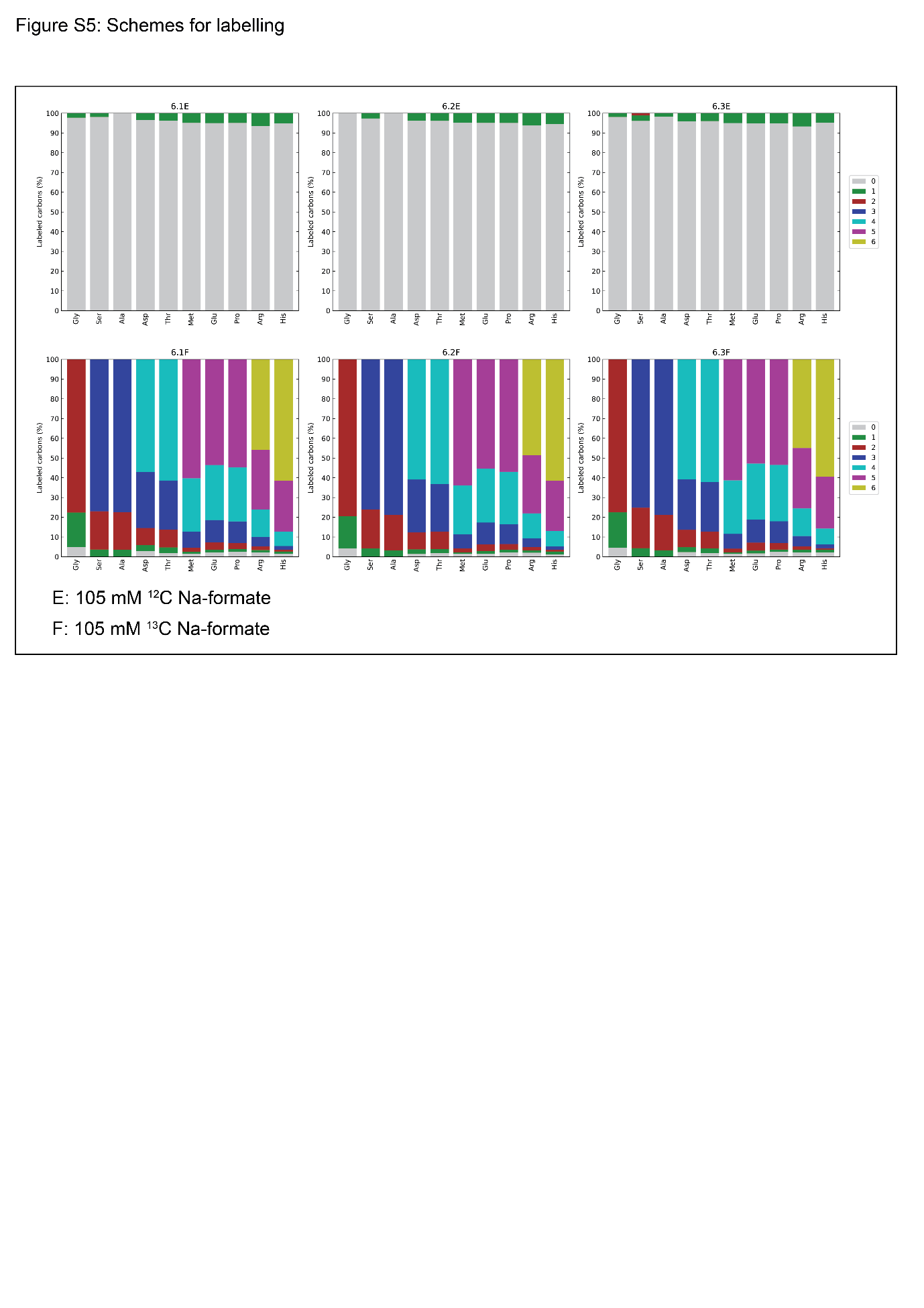

**Supplementary Figure 5: ^13^C isotopic labelling confirms full activity of the STC.**

The strain *STC2evo* in triplicates (6.1, 6.2 and 6.3) was cultivated in HMM200 supplemented with 105 mM ^12^C or ^13^C Na-formate and its incorporation into proteinogenic amino acids was analysed via liquid chromatography–mass spectrometry. (A) Labelling pattern of strains cultivated with ^12^C Na-formate and (B) Labelling pattern of strains cultivated with ^13^C Na-formate. The labelling pattern confirms the full activity of the STC as discussed in the main text. The experiment (in triplicates) was repeated three times, which showed highly similar labelling pattern.

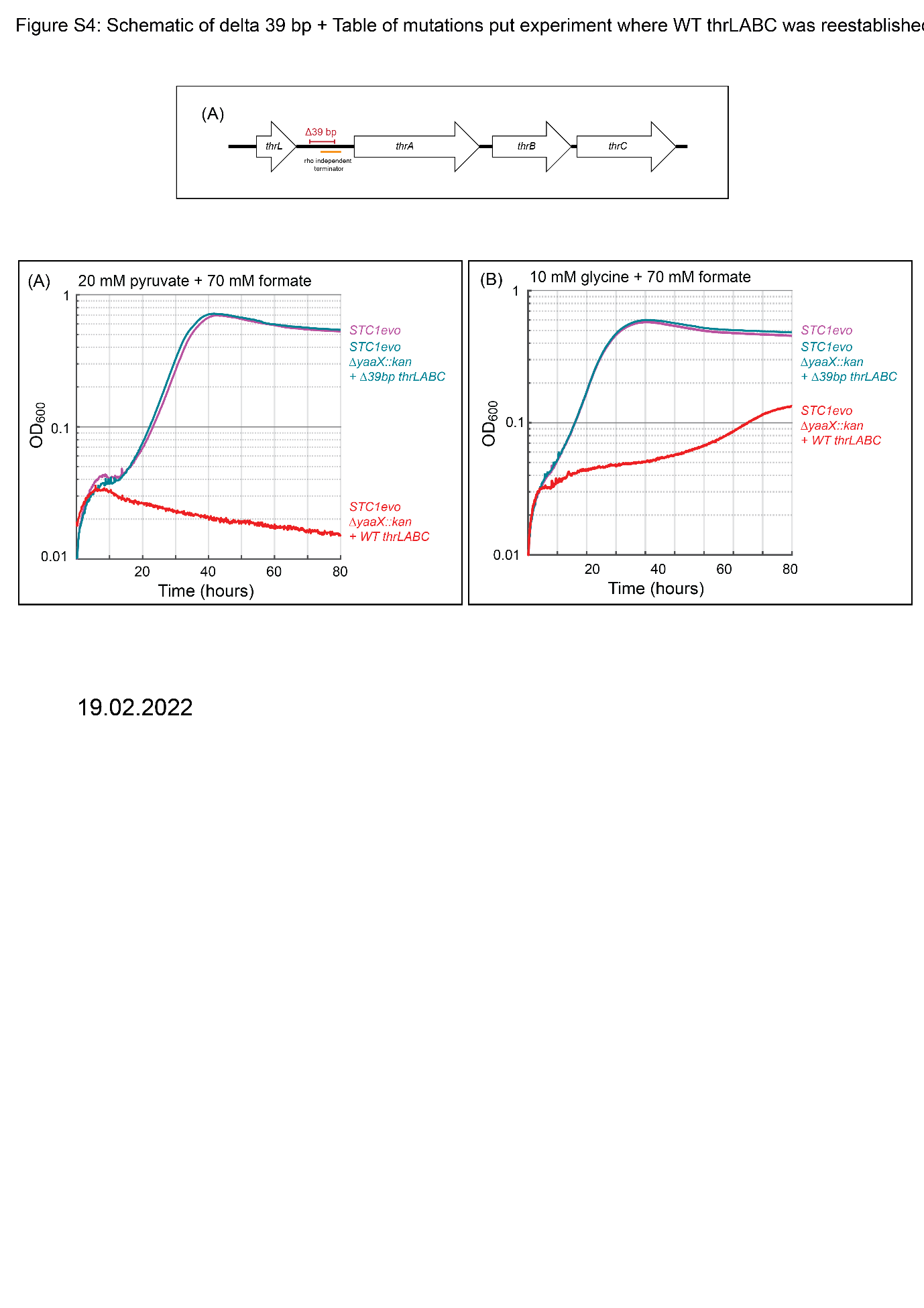

**Supplementary Figure 6: Reconstitution of WT module 4 abolishes growth via the STC.**

(A) In the ALE1 experiment a characteristic 39 bp deletion (Δ39 bp)emerged in the strains capable of growing on pyruvate and formate via the STC. The deletion removes a characteristic stem-loop located between the leader peptide thrL and the first gene of the threonine biosynthesis cluster thrA. The deletion likely abolishes attenuation leading to constitutive expression of the operon which equals M4 expression. To test the influence of Δ39 bp on the STC activity we reconstituted the WT *thrLABC* operon in the evolved strain (*STC1evo*) using P1 phage-mediated transfer and linkage (*yaaX* is a non-essential gene, located in close proximity to the *thrLABC* operon. Deleting the gene in a WT strain and transferring it to the *STC1evo* strain allowed us to obtain *STC1evoΔyaaX::kan* with and without the Δ39 bp mutation). (B) upon reconstitution of the WT *thrLABC* operon growth on pyruvate and formate is completely abolished and (C) growth on glycine and formate is severely impacted. All experiments (in triplicates) were repeated three times, which showed highly similar growth behavior.
