## Supplementary Tables for "Synthetic carbon fixation via the autocatalytic serine threonine cycle"

**Supplementary Table 1. Strains and plasmids used in this study**

| **Strain** | **Genotype** | **Source** | **Sequence location** |
| --- | --- | --- | --- |
| DH5α | *F^−^ λ^−^ Φ80lacZΔM15 Δ(lacZYA-argF)U169 deoR recA1 endA1 hsdR17(rK^−^ mK^+^) phoA supE44 thi-1 gyrA96 relA1* | ^1^ |  |
| ST18 | *pro thi hsdR^+^ Tp^r^ Sm^r^; chromosome::RP4-2 Tc::Mu-Kan::Tn7/λpir ΔhemA* | ^2^ |  |
| MG1655 | *F^−^ λ^−^ ilvG^−^ rfb-50 rph-1* | ^3^ |  |
| SIJ488 | MG1655 *Tn7::para-exo-beta-gam; prha-FLP; xylSpm-IsceI* | ^4^ |  |
| *XGAS* | SIJ488 Δ*aceA* Δ*aceE* Δ*pflB* Δ*poxB* Δ*serA* | This study |  |
| *XGAS:gM4gM5* | *XGAS*  P_STRONG_-RBS_C_ *thrABC,* P_STRONG_-RBS_C_ *kbl-tdh* | This study |  |
| *XGAS:gM5* | *XGAS*  P_STRONG_-RBS_C_ *kbl-tdh* | This study |  |
| *XGAS_C1:gM5* | *XGAS:gM5*  Δ*gcvTHP* | This study |  |
| *XGAS_C1:gM1gM5* | *XGAS_C1:gM5*  SS9: P_STRONG_ RBS_C_-*Me_ftfL-Me_fch-Me_mtdA* | This study |  |
| *XGAS_C1:gM1gM2gM3gM5eM (STC1)* | *XGAS_C1:gM1gM5*  P_STRONG_-RBS_B_ *glyA*  SS2: P_STRONG_ RBS_C_: *Cn_sdaA*  SS6: P_STRONG_ RBS_B_: *Re_pyc*  SS10: P_STRONG_ RBS_A_: *Ps_fdh* | This study | Genome Sequence Archive |
| *STC1evo* | *STC1* evolved in ALE1 with 20 mM pyruvate and 70 mM formate | This study | Genome Sequence Archive |
| *STC2* | *STC1evo*  *ΔiclR + aceA:aceB* | This study | Genome Sequence Archive |
| *STC2evo* | *STC2* evolved in ALE2 with limiting glycine and excess formate | This study | Genome Sequence Archive |

**Supplementary Table 2. Mutations detected by genome sequencing.**

Mutations that arose in ALE1 are marked in light yellow and mutations that arose in ALE2 are marked in dark yellow. Sequence analysis was performed using breseq^5^ with the STC1 genome as a reference.

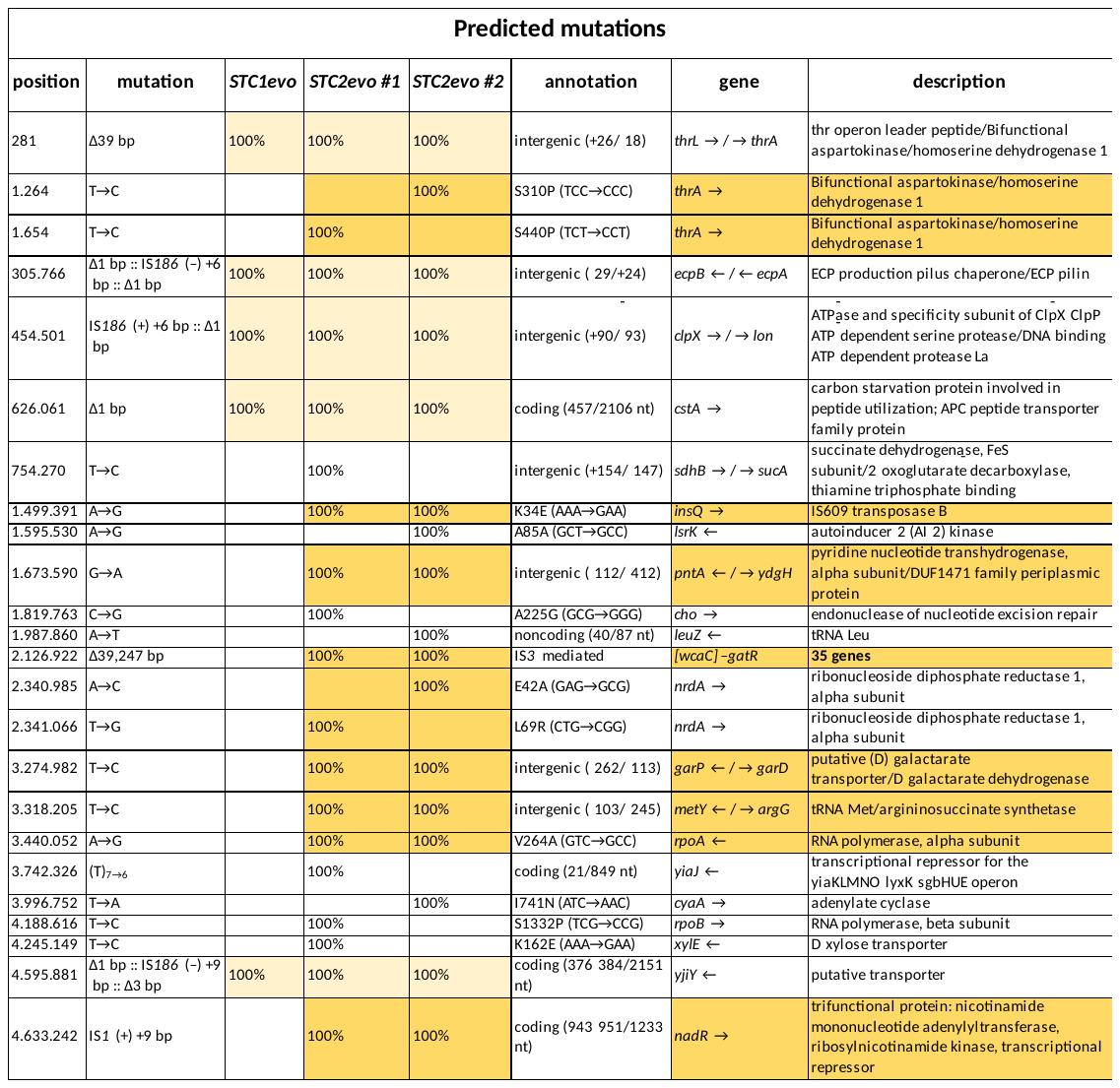
