## Supplementary material for "Synthetic carbon fixation via the autocatalytic serine threonine cycle": Gene Sequences

**Gene sequences as used in this study**

Formate-tetrahydrofolate ligase from *Methylobacterium extorquens* AM1 (UniProt: Q83WS0)

ATGCATCATCACCATCACCACCCGTCTGACATCGAGATCGCTCGTGCTGCTACCCTGAAGCCGATCGCTCAGGTTGCTGAGAAGCTGGGTATCCCGGACGAAGCTCTGCACAACTACGGTAAACACATCGCTAAAATCGACCACGACTTCATCGCTTCTCTGGAAGGTAAACCGGAAGGTAAACTGGTTCTGGTTACCGCTATCTCTCCGACCCCGGCTGGTGAAGGTAAAACCACCACCACCGTTGGTCTGGGTGACGCTCTGAACCGTATCGGTAAACGTGCTGTTATGTGCCTGCGTGAACCGTCTCTGGGTCCGTGCTTTGGTATGAAGGGTGGTGCTGCTGGTGGTGGTAAAGCTCAGGTTGTTCCGATGGAACAGATCAACCTGCACTTCACCGGTGACTTCCACGCTATCACCTCTGCTCACTCTCTGGCTGCTGCTCTGATCGACAACCACATCTACTGGGCTAACGAACTGAACATCGACGTGCGTCGTATCCACTGGCGTCGTGTTGTTGACATGAACGACCGTGCTCTGCGTGCTATCAACCAGTCTCTGGGTGGTGTTGCTAACGGTTTCCCGCGTGAAGACGGTTTTGACATCACCGTTGCTTCTGAGGTTATGGCTGTGTTCTGCCTGGCCAAAAACCTGGCTGACCTGGAAGAACGTCTGGGTCGTATCGTTATCGCTGAGACCCGTGACCGTAAACCGGTTACCCTGGCTGACGTTAAAGCTACCGGTGCTATGACCGTTCTGCTGAAGGACGCTCTGCAACCGAACCTGGTTCAGACCCTGGAAGGTAACCCGGCTCTGATCCACGGTGGTCCGTTTGCTAACATCGCTCACGGTTGCAACTCTGTTATCGCTACCCGTACCGGTCTGCGTCTGGCTGACTACACCGTTACCGAAGCTGGTTTTGGTGCTGACCTGGGTGCTGAGAAGTTCATCGACATCAAATGCCGTCAAACTGGTCTGAAGCCGTCTGCTGTTGTTATCGTTGCTACCATCCGTGCTCTGAAGATGCACGGTGGTGTTAACAAAAAAGACCTGCAAGCTGAGAACCTGGACGCTCTGGAGAAGGGTTTTGCTAACCTGGAACGTCACGTTAACAACGTGCGTTCTTTTGGTCTGCCGGTTGTTGTTGGTGTTAACCACTTCTTCCAGGACACCGACGCTGAACACGCTCGTCTGAAGGAACTGTGCCGTGACCGTCTGCAAGTTGAAGCTATCACCTGCAAACACTGGGCTGAAGGTGGTGCTGGTGCTGAAGCTCTGGCTCAGGCTGTTGTTAAACTGGCTGAAGGTGAACAGAAGCCGCTGACCTTTGCTTACGAGACCGAGACCAAAATCACCGACAAAATCAAAGCTATCGCTACCAAACTGTACGGTGCTGCTGACATCCAGATCGAATCTAAAGCTGCTACCAAACTGGCTGGTTTTGAGAAGGACGGTTACGGTGGCCTGCCGGTCTGCATGGCAAAAACCCAGTACTCTTTCTCTACCGACCCGACCCTGATGGGTGCTCCGTCTGGCCACCTGGTGAGCGTGCGTGACGTGCGTCTGTCTGCTGGTGCTGGTTTTGTTGTTGTTATCTGCGGTGAGATCATGACCATGCCGGGTCTGCCGAAGGTTCCGGCTGCTGACACCATCCGTCTGGACGCTAACGGTCAGATCGACGGTCTGTTCTAA

5,10-methenyltetrahydrofolate cyclohydrolase from *Methylobacterium extorquens* AM1 (UniProt: P55818)

ATGCATCATCACCATCACCACGCTGGTAACGAAACCATCGAAACCTTCCTGGACGGTCTGGCTTCTTCTGCTCCGACCCCGGGTGGTGGTGGTGCTGCTGCTATCTCTGGTGCTATGGGTGCTGCTCTGGTTTCTATGGTTTGCAACCTGACCATCGGTAAAAAAAAATACGTTGAAGTTGAAGCTGACCTGAAACAGGTTCTGGAAAAATCTGAAGGTCTGCGTCGTACCCTGACCGGTATGATCGCTGACGACGTTGAAGCTTTCGACGCTGTTATGGGTGCTTACGGTCTGCCGAAAAACACCGACGAAGAAAAAGCTGCTCGTGCTGCTAAAATCCAGGAAGCTCTGAAAACCGCTACCGACGTTCCGCTGGCTTGCTGCCGTGTTTGCCGTGAAGTTATCGACCTGGCTGAAATCGTTGCTGAAAAAGGTAACCTGAACGTTATCTCTGACGCTGGTGTTGCTGTTCTGTCTGCTTACGCTGGTCTGCGTTCTGCTGCTCTGAACGTTTACGTTAACGCTAAAGGTCTGGACGACCGTGCTTTCGCTGAAGAACGTCTGAAAGAACTGGAAGGTCTGCTGGCTGAAGCTGGTGCTCTGAACGAACGTATCTACGAAACCGTTAAATCTAAAGTTAACTAA

5,10-methylenetetrahydrofolate dehydrogenase from *Methylobacterium extorquens* AM1 (UniProt: P55818)

ATGCATCATCACCATCACCACTCTAAAAAACTGCTGTTCCAGTTCGACACCGACGCTACCCCGTCTGTTTTCGACGTTGTTGTTGGTTACGACGGTGGTGCTGACCACATCACCGGTTACGGTAACGTTACCCCGGACAACGTTGGTGCTTACGTTGACGGTACCATCTACACCCGTGGTGGTAAAGAAAAACAGTCTACCGCTATCTTCGTTGGTGGTGGTGACATGGCTGCTGGTGAACGTGTTTTCGAAGCTGTTAAAAAACGTTTCTTCGGTCCGTTCCGTGTTTCTTGCATGCTGGACTCTAACGGTTCTAACACCACCGCTGCTGCTGGTGTTGCTCTGGTTGTTAAAGCTGCTGGTGGTTCTGTTAAAGGTAAAAAAGCTGTTGTTCTGGCTGGTACCGGTCCGGTTGGTATGCGTTCTGCTGCTCTGCTGGCTGGTGAAGGTGCTGAAGTTGTTCTGTGCGGTCGTAAACTGGACAAAGCTCAGGCTGCTGCTGACTCTGTTAACAAACGTTTCAAAGTTAACGTTACCGCTGCTGAAACCGCTGACGACGCTTCTCGTGCTGAAGCTGTTAAAGGTGCTCACTTCGTTTTCACCGCTGGTGCTATCGGTCTGGAACTGCTGCCGCAGGCTGCTTGGCAGAACGAATCTTCTATCGAAATCGTTGCTGACTACAACGCTCAGCCGCCGCTGGGTATCGGTGGTATCGACGCTACCGACAAAGGTAAAGAATACGGTGGTAAACGTGCTTTCGGTGCTCTGGGTATCGGTGGTCTGAAACTGAAACTGCACCGTGCTTGCATCGCTAAACTGTTCGAATCTTCTGAAGGTGTTTTCGACGCTGAAGAAATCTACAAACTGGCTAAAGAAATGGCTTAA

Serine deaminase from *Cupriavidus necator* (UniProt: Q0K5P2)

ATGCATCATCACCATCACCACGTTGCTGTTTCTGTTTTCGACCTGTTCAAAGTTGGTATCGGTCCGTCTTCTTCTCACACCGTTGGTCCGATGCGTGCTGCTCTGATGTTCGCTCAGGGTCTGGAACGTGACGGTCTGCTGCCGCAGGTTGCTTCTGTTCGTGCTGAACTGTACGGTTCTCTGGGTGCTACCGGTAAAGGTCACGGTACCGACAAAGGTGTTATCCTGGGTCTGATGGGTGAAGCTCCGGACACCATCGACCCGGACTCTATCGACACCCGTCTGGCTGCTCTGCGTGCTGGTCGTGAACTGTCTCTGCTGGGTCAGCACGTTGTTCCGTTCGTTGAAAAAGAACACATCGCTTTCTACCGTCGTGAAGCTATGGCTGAACACCCGAACGGTATGAAATTCCACGCTTTCGACGCTGGTGGTGCTTCTCTGCGTGAAGCTCGTTACCTGTCTGTTGGTGGTGGTTTCGTTGTTACCGCTGGTGCTCCGAACACCCAGGTTCTGAACGCTGCTCAGCAGCTGCCGCACCCGTTCCGTTCTGGTAAAGACATGCTGGAAATGGCTACCGCTTCTGGTAAATCTATCGCTCGTCTGATGATGGAAAACGAACTGACCTGGCGTTCTGAAGCTGAAGTTACCTCTGGTCTGCTGCACATCTGGGACGTTATGCAGGCTTGCGTTGCTCGTGGTTGCCGTACCGACGGTGAACTGCCGGGTCCGTTCCGTGTTAAACGTCGTGCTCCGGAACTGTTCCGTTCTCTGACCGAACGTGCTGAACGTACCCTGTCTGACCCGCTGTCTGTTATGGACTGGGTTAACCTGTACGCTATCGCTGTTAACGAAGAAAACGCTGCTGGTGGTCGTGTTGTTACCGCTCCGACCAACGGTGCTGCTGGTATCATCCCGGCTGTTCTGCACTACTACGACCGTTTCGTTCCGGGTGCTTCTAAACAGGGTGTTGTTGACTTCCTGCTGACCGCTGGTGCTATCGGTCTGCTGTACAAACTGAACGCTTCTATCTCTGGTGCTGAAGTTGGTTGCCAGGGTGAAGTTGGTGTTGCTTGCTCTATGGCTGCTGGTGCTCTGGCTGCTGTTCAGGGTGGTTCTCCGGCTCAGGTTGAAAACGCTGCTGAAATCGGTATGGAACACAACCTGGGTCTGACCTGCGACCCGGTTGGTGGTCTGGTTCAGATCCCGTGCATCGAACGTAACGCTATGGCTTCTGTTAAAGCTGTTAACGCTGCTCGTATGGCTCTGCGTGGTGACGGTACCCACTACGTTTCTCTGGACTCTGTTATCAAAACCATGCGTGAAACCGGTGCTGACATGAAAACCAAATACAAAGAAACCGCTCGTGGTGGTCTGGCTGTTAACATCGTTGAATGCTAA

Pyruvate carboxylase from *Rhizobium eetli* (UniProt: Q2K340)

TCATCACCATCACCACCCGATCTCTAAAATCCTGGTTGCTAACCGTTCTGAAATCGCTATCCGTGTTTTCCGTGCTGCTAACGAACTGGGTATCAAAACCGTTGCTATCTGGGCTGAAGAAGACAAACTGGCTCTGCACCGTTTCAAAGCTGACGAATCTTACCAGGTTGGTCGTGGTCCGCACCTGGCTCGTGACCTGGGTCCGATAGAATCTTACCTGTCTATCGACGAAGTTATCCGTGTTGCTAAACTGTCTGGTGCTGACGCTATCCACCCGGGTTACGGTCTGCTGTCTGAATCTCCGGAATTTGTTGACGCTTGCAACAAAGCTGGTATCATCTTCATCGGTCCGAAAGCTGACACCATGCGTCAGCTGGGTAACAAAGTTGCTGCTCGTAACCTGGCTATCTCTGTTGGTGTTCCGGTTGTTCCGGCTACCGAACCGCTGCCGGACGACATGGCTGAAGTTGCTAAAATGGCTGCTGCTATCGGTTACCCGGTTATGCTGAAAGCTTCTTGGGGTGGTGGTGGTCGTGGTATGCGTGTTATCCGTTCTGAAGCTGACCTGGCTAAAGAAGTTACCGAAGCTAAACGTGAAGCTATGGCTGCTTTCGGTAAAGACGAAGTTTACCTGGAAAAACTGGTTGAACGTGCTCGTCACGTTGAATCTCAGATCCTGGGTGACACCCACGGTAACGTTGTTCACCTGTTCGAACGTGACTGCTCTGTTCAGCGTCGTAACCAGAAAGTTGTTGAACGTGCTCCGGCTCCGTACCTGTCTGAAGCTCAGCGTCAGGAACTGGCTGCTTACTCTCTGAAAATCGCTGGTGCTACCAACTACATCGGTGCTGGTACCGTTGAATACCTGATGGACGCTGACACCGGTAAATTCTACTTCATCGAAGTTAACCCGCGTATCCAGGTTGAACACACCGTTACCGAAGTTGTTACCGGTATCGACATCGTTAAAGCTCAGATCCACATCCTGGACGGTGCTGCTATCGGTACCCCGCAGTCTGGTGTTCCGAACCAGGAAGACATCCGTCTGAACGGTCACGCTCTGCAATGCCGTGTTACCACCGAAGACCCGGAACACAACTTCATCCCGGACTACGGTCGTATCACCGCTTACCGTTCTGCTTCTGGTTTCGGTATCCGTCTGGACGGTGGTACCTCTTACTCTGGTGCTATCATCACCCGTTACTACGACCCGCTGCTGGTTAAAGTTACCGCTTGGGCTCCGAACCCGCTGGAAGCTATCTCTCGTATGGACCGTGCTCTGCGTGAATTTCGTATCCGTGGTGTTGCTACCAACCTGACCTTCCTGGAAGCTATCATCGGTCACCCGAAATTCCGTGACAACTCTTACACCACCCGTTTCATCGACACCACCCCGGAACTGTTCCAGCAGGTTAAACGTCAGGACCGTGCTACCAAACTGCTGACCTACCTGGCTGACGTTACCGTTAACGGTCACCCGGAAGCTAAAGACCGTCCGAAACCGCTGGAAAACGCTGCTCGTCCGGTTGTTCCGTACGCTAACGGTAACGGTGTTAAAGACGGTACCAAACAGCTGCTGGACACCCTGGGTCCGAAAAAATTCGGTGAATGGATGCGTAACGAAAAACGTGTTCTGCTGACCGACACCACCATGCGTGACGGTCACCAGTCTCTGCTGGCTACCCGTATGCGTACCTACGACATCGCTCGTATCGCTGGTACCTACTCTCACGCTCTGCCGAACCTGCTGTCTCTGGAATGCTGGGGTGGTGCTACCTTCGACGTTTCTATGCGTTTCCTGACCGAAGACCCGTGGGAACGTCTGGCTCTGATCCGTGAAGGTGCTCCGAACCTGCTGCTGCAAATGCTGCTGCGTGGTGCTAACGGTGTTGGTTACACCAACTACCCGGACAACGTTGTTAAATACTTCGTTCGTCAGGCTGCTAAAGGTGGTATCGACCTGTTCCGTGTTTTCGACTGCCTGAACTGGGTTGAAAACATGCGTGTTTCTATGGACGCTATCGCTGAAGAAAACAAACTGTGCGAAGCTGCTATCTGCTACACCGGTGACATCCTGAACTCTGCTCGTCCGAAATACGACCTGAAATACTACACCAACCTGGCTGTTGAACTGGAAAAAGCTGGTGCTCACATCATCGCTGTTAAAGACATGGCTGGTCTGCTGAAACCGGCTGCTGCTAAAGTTCTGTTCAAAGCTCTGCGTGAAGCTACCGGTCTGCCGATCCACTTCCACACCCACGACACCTCTGGTATCGCTGCTGCTACCGTTCTGGCTGCTGTTGAAGCTGGTGTTGACGCTGTTGACGCTGCTATGGACGCTCTGTCTGGTAACACCTCTCAGCCGTGCCTGGGTTCTATCGTTGAAGCTCTGTCTGGTTCTGAACGTGACCCGGGTCTGGACCCGGCTTGGATCCGTCGTATCTCTTTCTACTGGGAAGCTGTTCGTAACCAGTACGCTGCTTTCGAATCTGACCTGAAAGGTCCGGCTTCTGAAGTTTACCTGCACGAAATGCCGGGTGGTCAGTTCACCAACCTGAAAGAACAGGCTCGTTCTCTGGGTCTGGAAACCCGTTGGCACCAGGTTGCTCAGGCTTACGCTGACGCTAACCAGATGTTCGGTGACATCGTTAAAGTTACCCCGTCTTCTAAAGTTGTTGGTGACATGGCTCTGATGATGGTTTCTCAGGACCTGACCGTTGCTGACGTTGTTTCTCCGGACCGTGAAGTTTCTTTCCCGGAATCTGTTGTTTCTATGCTGAAAGGTGACCTGGGTCAGCCGCCGTCTGGTTGGCCGGAAGCTCTGCAAAAAAAAGCTCTGAAAGGTGAAAAACCGTACACCGTTCGTCCGGGTTCTCTGCTGAAAGAAGCTGACCTGGACGCTGAACGTAAAGTTATCGAAAAAAAACTGGAACGTGAAGTTTCTGACTTCGAATTTGCTTCTTACCTGATGTACCCGAAAGTTTTCACCGACTTCGCTCTGGCTTCTGACACCTACGGTCCGGTTTCTGTTCTGCCGACCCCGGCTTACTTCTACGGTCTGGCTGACGGTGAAGAACTGTTCGCTGACATCGAAAAAGGTAAAACCCTGGTTATCGTTAACCAGGCTGTTTCTGCTACCGACTCTCAGGGTATGGTTACCGTTTTCTTCGAACTGAACGGTCAGCCGCGTCGTATCAAAGTTCCGGACCGTGCTCACGGTGCTACCGGTGCTGCTGTTCGTCGTAAAGCTGAACCGGGTAACGCTGCTCACGTTGGTGCTCCGATGCCGGGTGTTATCTCTCGTGTTTTCGTTTCTTCTGGTCAGGCTGTTAACGCTGGTGACGTTCTGGTTTCTATCGAAGCTATGAAAATGGAAACCGCTATCCACGCTGAAAAAGACGGTACCATCGCTGAAGTTCTGGTTAAAGCTGGTGACCAGATCGACGCTAAAGACCTGCTGGCTGTTTACGGTGGTTAA

Formate dehydrogenase from *Pseudomonas* sp. 101 (UniProt: P33160)

ATGCATCATCACCATCACCACGCTAAAGTTCTGTGCGTTCTGTACGACGACCCGGTTGACGGTTACCCGAAAACCTACGCTCGTGACGACCTGCCGAAAATCGACCACTACCCGGGTGGTCAGACCCTGCCGACCCCGAAAGCTATCGACTTCACCCCGGGTCAGCTGCTGGGTTCTGTTTCTGGTGAACTGGGTCTGCGTAAATACCTGGAATCTAACGGTCACACCCTGGTTGTTACCTCTGACAAAGACGGTCCGGACTCTGTTTTCGAACGTGAACTGGTTGACGCTGACGTTGTTATCTCTCAGCCGTTCTGGCCGGCTTACCTGACCCCGGAACGTATCGCTAAAGCTAAAAACCTGAAACTGGCTCTGACCGCTGGTATCGGTTCTGACCACGTTGACCTGCAATCTGCTATCGACCGTAACGTTACCGTTGCTGAAGTTACCTACTGCAACTCTATCTCTGTTGCTGAACACGTTGTTATGATGATCCTGTCTCTGGTTCGTAACTACCTGCCGTCTCACGAATGGGCTCGTAAAGGTGGTTGGAACATAGCTGACTGCGTAAGCCACGCTTACGACCTGGAAGCTATGCACGTTGGTACCGTTGCTGCTGGTCGTATCGGTCTGGCTGTTCTGCGTCGTCTGGCTCCGTTCGACGTTCACCTGCACTACACCGACCGTCACCGTCTGCCGGAATCTGTTGAAAAAGAACTGAACCTGACCTGGCACGCTACCCGTGAAGACATGTACCCGGTTTGCGACGTTGTTACCCTGAACTGCCCGCTGCACCCGGAAACCGAACACATGATCAACGACGAAACCCTGAAACTGTTCAAACGTGGTGCTTACATCGTTAACACCGCTCGTGGTAAACTGTGCGACCGTGACGCTGTTGCTCGTGCTCTGGAATCTGGTCGTCTGGCTGGTTATGCGGGTGACGTGTGGTTCCCCCAGCCGGCTCCGAAAGACCACCCGTGGCGTACCATGCCGTACAACGGTATGACCCCGCACATCTCTGGTACCACCCTGACCGCTCAGGCTCGTTACGCTGCTGGTACCCGTGAAATCCTGGAATGCTTCTTCGAAGGTCGTCCGATCCGTGACGAATACCTGATCGTTCAGGGTGGTGCTCTGGCTGGTACCGGTGCTCACTCTTACTCTAAAGGTAACGCTACCGGTGGTTCTGAAGAAGCTGCTAAATTCAAAAAAGCTGTTTAA
